# Acid phospholipase A1 promotes lysosomal membrane catabolism

**DOI:** 10.64898/2026.08.07.743472

**Authors:** Martin Tischitz, Johannes Breithofer, Dominik Bulfon, Clara Zitta, Amit Singh Sahrawat, Carina Wagner, Nermeen Fawzy, Thomas Züllig, Monika Oberer, Lennart Hartig, Anita Pirchheim, Laszlo Schooltink, Ulrike Taschler, Karl Gruber, Achim Lass, Ulrich Stelzl, Dagmar Kolb, Dagmar Kratky, Robert Zimmermann

## Abstract

The molecular mechanisms of lysosomal glycerophospholipid (GPL) catabolism are incompletely understood. Here, we report that acid phospholipase A1 (APLA1), formerly known as palmitoyl-protein thioesterase 2 (PPT2), is required for efficient GPL degradation. Deletion of *APLA1* in human cells results in excess accumulation of phospholipids within lysosomes # a pathological condition termed phospholipidosis. APLA1 activity depends on interactions with negatively charged GPLs and is inhibited by phospholipidosis-inducing cationic amphiphilic drugs. Hydrolysis of zwitterionic, but not anionic, GPLs requires co-activation of APLA1 by the lysosome-specific lipid bis(monoacylglycero)phosphate. Upon pharmacological mTORC inhibition, which increases lysosomal GPL turnover, *APLA1*-deficient cells exhibit massive accumulation of multilamellar membranes in lysosomes and reduced cytosolic triacylglycerol stores. APLA1 acts in concert with lysosomal phospholipase A2 (PLA2G15). Combined *APLA1/PLA2G15*-deficiency leads to a severe reduction in acid phospholipase A1/A2 activity, thereby exacerbating phospholipidosis. Our observations provide detailed mechanistic insights into lysosomal GPL catabolism, a crucial pathway for maintaining lipid homeostasis.

## Introduction

Lysosomes internalize biomembranes through endocytic or autophagic pathways for catabolism and recycling. This process is essential for maintaining cellular lipid and energy homeostasis. Disturbances in lysosomal catabolism of sphingolipids, neutral lipids, and glycerophospholipids (GPL) are the underlying cause of numerous lysosomal storage disorders and are frequently associated with neurodegeneration^1,2^.

GPLs are the predominant lipid species of membranes. Their hydrolysis in lysosomes requires phospholipases which are active in the acidic lumen of these organelles. According to our current knowledge, GPL catabolism is mediated by acid phospholipase A1 and A2 activity (PLA1 and PLA2), cleaving fatty acid (FA) ester bonds at the *sn-1* and *sn-2* position, respectively. The complete deacylation of GPLs produces FAs and glycerophosphodiesters (GPDs), which are exported from lysosomes for subsequent metabolism^3^. Acid GPL hydrolysis can be catalyzed by lysosomal phospholipase A2 group XV (PLA2G15) and phospholipase B domain containing 2 (PLBD2). PLA2G15 has long been considered the major lysosomal PLA1 and PLA2. It exhibits broad substrate specificity for anionic and zwitterionic GPLs^4^, while PLBD2 selectively cleaves inositol-containing lyso-GPLs^5^. However, accumulating evidence indicates that PLA2G15 alone cannot account for lysosomal GPL catabolism, as *PLA2G15*-deficiency causes only moderate changes in lysosomal GPL content and composition^6,7^. These observations strongly suggest the existence of additional lysosomal acid phospholipases.

As an alternative to PLA1/2-catalyzed mechanisms, GPL degradation may be initiated by the removal of polar head groups by phospholipase C (PLC), resulting in the release of diacylglycerol (DAG) and phosphorylated headgroups. The GPL-derived DAG can be further hydrolyzed into FAs and glycerol by lysosomal acid lipase (LAL), the major acylglycerol and cholesteryl ester hydrolase in lysosomes^8^. Acid sphingomyelinase (ASM) is the only known lysosomal PLC and has been reported to hydrolyze sphingomyelins, but also GPLs, lyso-GPLs, and plasmalogens in vitro^9^. However, *ASM*-deficiency is primarily characterized by the excessive accumulation of sphingolipids^10^, arguing against a major role in GPL catabolism.

Together, the available data indicate that lysosomal GPL degradation cannot be fully explained by previously identified acid phospholipases. Here, we identify palmitoyl protein thioesterase 2 (PPT2) as the long-sought acid phospholipase. Together with PLA2G15, it forms the core of the lysosomal GPL degradation machinery.

## Results

### PPT2 acts as acid phospholipase A1

To identify new lysosomal phospholipases A1/A2, we performed an unbiased screen of ∼200 murine enzymes for phospholipase activity under acidic conditions (pH 4.5). The enzyme library included the majority of known lipid hydrolases as well as structurally related enzymes of unknown function^11^. We transiently expressed these enzymes in Expi293F cells and used cell lysates as source of enzymatic activity. Substrate hydrolysis was monitored by the detection of liberated FAs using a colorimetric assay (**Fig. 1A**).

**Figure 1:**
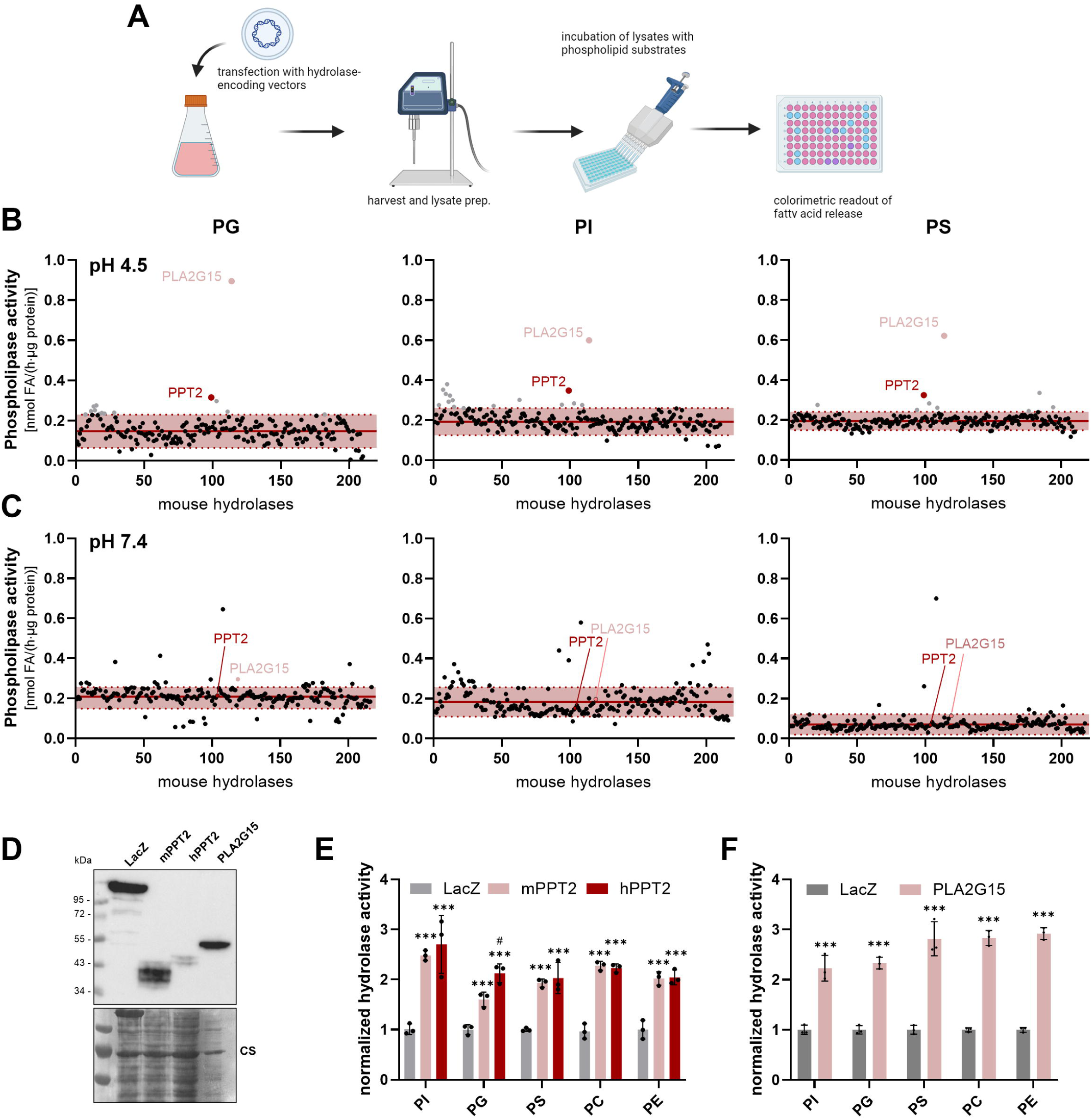
Mouse and human PPT2 exhibit acid phospholipase activity. **(A)** Schematic illustration of the screening workflow. Cell lysates of enzyme-expressing Expi293F cells (2 mg protein/ml) were incubated with 1 mM lipid substrate solution containing 2% BSA at acidic (4.5) or neutral (7.4) pH for 1 h at 37°C. FA release was then determined using a colorimetric assay. **(B)** Screening of murine lipid hydrolases for activity against phosphatidylglycerol (PG), phosphatidylinositol (PI), and phosphatidylserine (PS) at pH 4.5 and (**C**) pH 7.4. Red lines and red dotted lines indicate the overall mean ± SD of all data points. Enzymes with activities exceeding the overall mean + SD were considered positive hits. Red dots indicate lysosomal hydrolases and grey dots indicate positive hits not localizing to lysosomes (n = 1 for each condition). **(D)** Western blot analysis of Expi293F cells expressing HIS-tagged β-galactosidase (LacZ), mouse and human PPT2 (mPPT2 and hPPT2), and human PLA2G15. (**E** and **F**) Comparison of the head group specificity of PPT2 orthologues and PLA2G15. Lysates of Expi293F cells expressing mPPT2, hPPT2, PLA2G15, or LacZ were incubated with the indicated lipid substrates and the release of FAs was monitored (n = 3 independent replicates). Data are represented as mean ± SD. Statistically significant differences were determined by two-way ANOVA for (E) and (F) and corrected for multiple comparisons by Bonferroni post-hoc test (*: compared to LacZ, ^#^: compared to mPPT2, *^,#^*p*<0.05, \*\*\**p*<0.001)

Under the applied conditions, we identified several candidates capable of hydrolyzing the anionic GPLs phosphatidylglycerol (PG), phosphatidylinositol (PI), and phosphatidylserine (PS) (**Fig. 1B**). Among these candidates, only PLA2G15^4^ and PPT2^12^ localize to lysosomes. The other identified enzymes are presumably active in extra-lysosomal compartments and were therefore not investigated further. In line with the lysosomal localization, lysates of PPT2# and PLA2G15-expressing cells did not show increased activity at pH 7.4 (**Fig. 1C**). PPT2 is a poorly characterized lysosomal enzyme with acyl-CoA thioesterase activity^12^. To further investigate the role of the enzyme in lysosomal GPL metabolism, we compared the activities of mouse and human orthologues against a set of GPL substrates, again using lysates of enzyme-expressing cells as source of activity (**Fig. 1D**). We found that recombinant expression of both orthologues significantly increased acid phospholipase activity in comparison to the β-galactosidase-expressing control (LacZ). The enzymes hydrolyzed anionic substrates (PG, PI, PS) as well as the zwitterionic GPLs phosphatidylcholine and phosphatidylethanolamine (PC and PE) (**Fig. 1E**). Similar activities were observed for the positive control PLA2G15 (**Fig. 1F**). For reasons that will become clear later, we hereafter refer to PPT2 as acid phospholipase A1 (APLA1).

For a more comprehensive analysis of APLA1 function, we expressed the HIS-tagged human enzyme and a variant lacking the active serine (S111A) in Expi293F cells. APLA1 was partially secreted from cells, which is frequently observed for lysosomal proteins. APLA1 and its inactive variant were purified from supernatants by affinity chromatography, resulting in the enrichment of a protein fraction with a molecular weight ranging from ∼ 45 to 60 kDa (**Fig. S1A, B**). Deglycosylation by PNGase F resulted in a single band at the expected MW of 34 kDa, suggesting that APLA1 is heavily glycosylated (**Fig 2A**, Inset). Using PG as substrate, purified APLA1 exhibited specific activities between 10 and 20 nmol FA/(h·µg protein), while the S111A mutant enzyme was inactive (**Fig. 2A**). PG hydrolysis increased linearly up to 30 min and up to an enzyme concentration of 0.8 µg/reaction (**Fig. S1C, D**).

**Figure 2:**
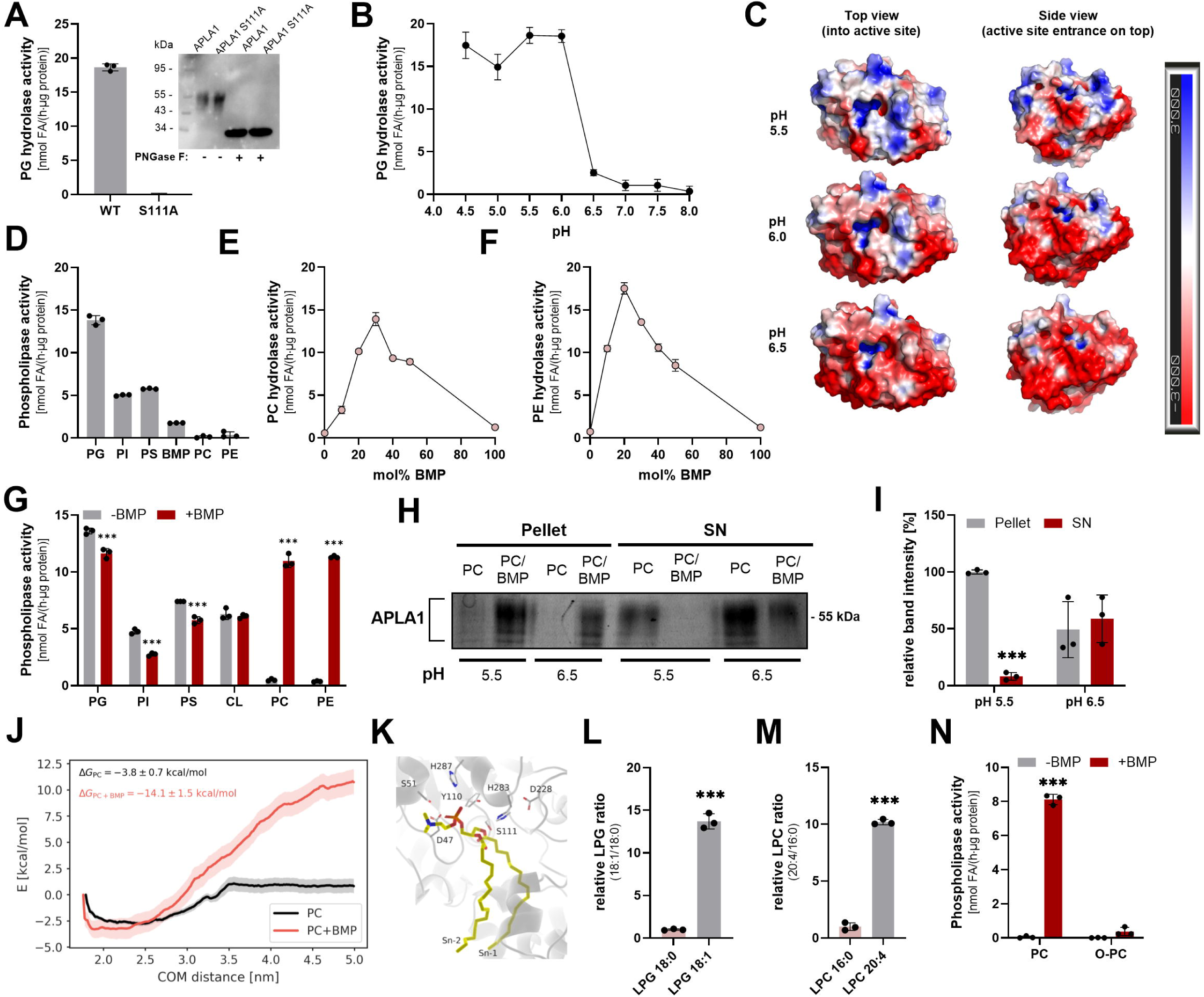
PPT2 acts as acid phospholipase A1 (APLA1). All activity assays were conducted with purified HIS-tagged APLA1. Lipid substrates were esterified with oleic acid, with the exception of PI (from bovine liver). Activities were determined by measuring the release of FAs using a colorimetric assay. Measurements were carried out at pH 4.5 for 30 min, unless otherwise stated, in the presence of 0.4 mM phospholipid substrates and 10 µg/ml APLA1 protein in a volume of 50 µl at 37°C. **(A)** Specific PG hydrolase activity of APLA1 and a mutant variant lacking the active serine (S111A) (n = 3 independent replicates, representative of 3 independent experiments). Inset: Western blot analysis of HIS-tagged APLA1 before and after deglycosylation with PNGase F. **(B)** pH dependence of APLA1 using PG as substrate (n = 3 independent replicates, data representative of three independent experiments). (**C**) Surface-charge-predictions for the solved crystal structure of human APLA1 (PDB-ID: 1PJA) at the indicated pH values. Blue and red areas indicate a positive and negative surface charge, respectively. (**D**) Head group specificity of APLA1 (n = 3 independent replicates, data are representative of two independent experiments). (**E** and **F**) Dose-dependent stimulation of APLA1-mediated PC and PE hydrolysis by *sn*-3,3’# S,S#bis(monoacylglycerol)phosphate (BMP) (n = 3 independent replicates, data are representative of two independent experiments). **(G)** APLA1-mediated hydrolysis of acidic (PG, PI, PS, and cardiolipin (CL)) and zwitterionic (PC and PE) GPLs. Phospholipase activity was detected in the absence or presence of 30 mol% BMP (n = 3 independent replicates, data representative of two independent experiments). (**H, I**) Binding of APLA1 to liposomes consisting either of PC or a mixture of PC/BMP (70/30 mol%). Liposomes were incubated with purified APLA1 at the indicated pH for 10 min and subsequently pelleted by centrifugation (100,000 x g). The protein content of pellet and supernatant was analyzed by SDS-PAGE and Coomassie staining (n = 3 independent experiments). (**J**) Free energies of APLA1 (PDB-ID: 1PJA) binding to liposomes composed of PC 18:1-18:1 or a mix of PC and BMP (1:1). Membrane binding behavior was calculated by molecular dynamic simulations (COM: center of mass). (**K**) Deep-learning-assisted co-modeling of PC 18:1-18:1 in the active site of APLA1 (PDB-ID: 1PJA). (**L-M**) Positional preference of APLA1. The enzyme was incubated with (L) PG 18:0-18:1 and (M) PC 16:0-20:4 and BMP (30 mol%). The generated lyso-GPLs were monitored by targeted MS analysis (n = 3 independent replicates). (**N**) Purified APLA1 was incubated with PC 18:1-18:1 or ether-PC 1-O-16:0-2-18:1 (O-PC) in the absence and presence of BMP (30 mol%, n = 3 independent replicates). Data are represented as mean ± SD. Statistically significant differences were determined by Student’s two-tailed unpaired *t*-test (L, M) and two-way ANOVA with Sidák post hoc test (G, I, N) (\*\*\**p*<0.001).

Saturation kinetics revealed an apparent K_m_ value of 60 µM and a V_max_ of 12 nmol FA/(h·µg protein) (**Fig. S1E**). Consistent with its lysosomal localization, hydrolytic activity was highest between pH 4.5 and 6.0 and was almost completely abolished at pH values ≥6.5 (**Fig. 2B**). To understand the drop in APLA1 activity at pH 6.5, we calculated the surface charge of the enzyme at different pH values, based on the solved crystal structure^13^ (**Fig. 2C**). At pH 5.5 and 6.0, the areas around the active center are primarily positively charged, likely promoting the interaction with the anionic phospholipid substrate. The charge of these areas is almost completely reversed at pH 6.5, suggesting that surface charge interactions are the driving force mediating the interaction between APLA1 and its substrate.

To determine the headgroup specificity of purified APLA1, we performed activity assays with different di-oleoyl GPL species. APLA1 readily hydrolyzed the anionic substrates PG, PI, and PS, and also showed low activity against the lysosome-specific lipid bis(monoacylglycero)phosphate (BMP, *sn*-3,3’-S,S-BMP) (**Fig. 2D**). In contrast to what has been observed with lysates (**Fig. 1E**), purified APLA1 exhibited no activity against the zwitterionic substrates PC and PE, indicating that the purified enzyme requires a co-factor for the hydrolysis of zwitterionic phospholipids present in cell lysates, but not in purified enzyme preparations.

### APLA1 requires BMP as co-factor for the hydrolysis of zwitterionic GPLs

Intralumenal vesicles of acidic organelles exhibit a negative surface potential, enabling the interaction with positively charged lumenal proteins. This negative charge is primarily provided by the anionic GPL BMP, which serves as a cofactor for many lysosomal hydrolases^14^. Importantly, addition of BMP to the reactions enabled APLA1-mediated PC and PE degradation with maximal activities observed between 20 and 40 mol% BMP (**Fig. 2E, F**). In comparison, a substrate containing only BMP (100 %) was poorly degraded. Analysis of lyso-GPL formation by TLC revealed that the reactions with BMP-containing substrates generate LPC and LPE, but not LPG, which would be expected from BMP hydrolysis (**Fig. S1F-I**). In contrast to PC and PE, addition of BMP to the substrate did not increase the hydrolysis of negatively charged PG, PI, PS, and cardiolipin (CL), but instead moderately reduced their turnover (**Fig. 2G**).

To directly investigate the interaction of APLA1 with its substrate, we incubated the enzyme with liposomes composed of PC or a mix of PC and BMP (molar ratio 70/30) at pH 5.5 and 6.5. Subsequently, liposomes were pelleted by centrifugation, and both the pellet and supernatant fractions were analyzed by SDS-PAGE (**Fig. 2H**). When liposomes were composed solely of PC, APLA1 was almost exclusively detected in the supernatant fraction regardless of the pH value. Conversely, when liposomes were composed of PC/BMP, APLA1 was found in the pellet fraction at pH 5.5. At pH 6.5, the protein was distributed between pellet and supernatant, suggesting that BMP is indeed required for efficient enzyme-substrate interactions (**Fig. 2H, I**). Accordingly, molecular dynamics (MD) simulations of APLA1 binding to PC and PC/BMP bilayers indicated that APLA1 exhibits a strong preference for binding to PC/BMP over PC membranes (**Fig. 2J, S2A, B**).

To investigate the regiospecificity of APLA1, we first conducted a deep-learning-assisted co-folding of APLA1 with PC, PE, or PG as substrates, suggesting a higher preference for the *sn-1* acyl carbon over the *sn-2* acyl carbon near the active site serine (S111) in the catalytic center for all substrates (**Fig. 2K, S2C**). To validate these results, we incubated purified APLA1 with 1-stearoyl-2-oleoyl PG (PG 18:0-18:1) and 1-palmitoyl-2-arachidonyl PC (PC 16:0-20:4/BMP, 70/30 mol%). Analysis of the generated lyso-GPLs by LC-MS revealed that the enzyme predominantly produced LPG 18:1 and LPC 20:4, suggesting preferential hydrolysis of substrates at *sn-1* positions (**Fig. 2L, M**). In line, the *sn-1* alkyl ether of PC (O-PC) was barely hydrolyzed, even in the presence of BMP (**Fig. 2N**). Overall, these observations suggest that the enzyme acts as lysosomal phospholipase A1.

### Deletion of *APLA1* leads to phospholipidosis

Our observations indicate that APLA1 degrades GPLs in acidic compartments, possibly in concert with PLA2G15. To compare the role of these phospholipases in lysosomal lipid metabolism, we genetically deleted APLA1 (*APLA1-*KO), PLA2G15 (*PLA2G15-*KO), or both APLA1 and PLA2G15 (dKO) in HEK293 cells using CRISPR/Cas9 gene editing. Sanger sequencing confirmed the generation of homozygous knockout lines, and proteomic analysis revealed complete loss of APLA1, PLA2G15, or both phospholipases (**Fig. S3A, B)**.

Lysosomal storage disorders are characterized by the accumulation of undigested material in acidic compartments, leading to an increased number and/or size of acidic organelles. To assess the effect of phospholipase-deficiency on lysosomal morphology, we stained acidic compartments with LysoTracker and determined the average LysoTracker-positive (LT^+^) area per cell. *APLA1*-KO cells displayed a 4-fold increase in LT^+^ area compared to WT cells, while those of *PLA2G15*-KO cells were 2.1-fold elevated. dKO cells exhibited a prominent increase in LT^+^ areas, exceeding WT levels more than 10-fold, indicating that combined deletion of APLA1 and PLA2G15 has synergistic effects (**Fig. 3A, B**).

**Fig. 3:**
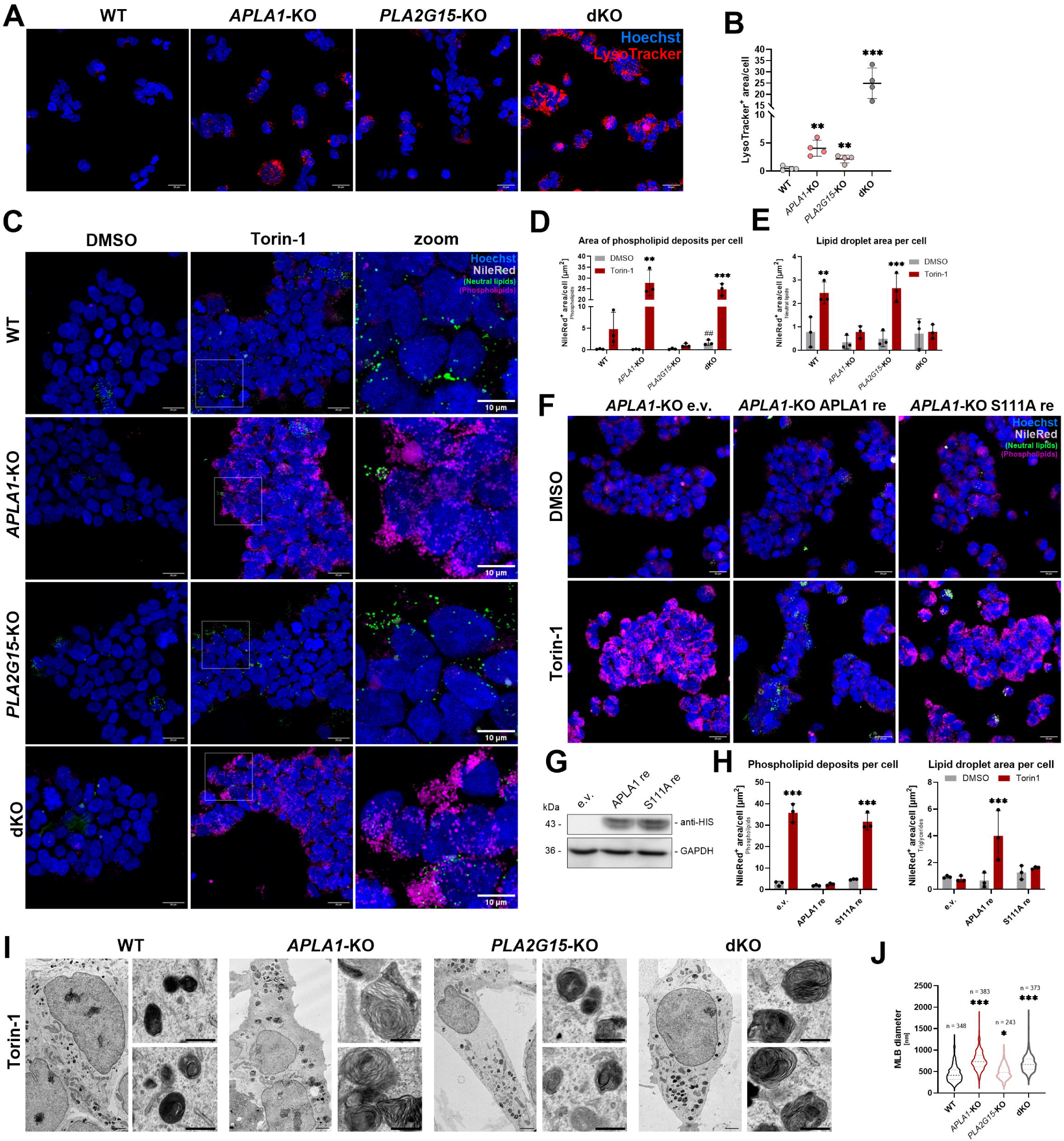
Loss of APLA1 in HEK 293 cells causes accumulation of phospholipids in lysosomes. (**A**) LysoTracker (red) staining of HEK293 WT, *APLA1*-KO, *PLA2G15*-KO, and dKO cells. Nuclei were stained with Hoechst (blue); scale bars: 20 µm. (**B**) Quantitative analysis of images acquired in (A) (n = 4 fields, data representative of two independent experiments). (**C**) NileRed staining for neutral lipids (green) and phospholipids (magenta) of cells treated with DMSO (vehicle) or 250 nM Torin-1 for 16 h. Nuclei were stained with Hoechst (blue); scale bars: 20 µm (overview), 10 µm (zoom). (**D, E**) Quantitative analysis of NileRed stained phospholipid deposits and LDs from (C) (n = 3 fields, data representative of two independent experiments). (**F**) Stable re-expression of APLA1 (APLA1 re) and the inactive APLA1 variant S111A (S111A re) in *APLA1*# KO cells. An empty vector was used as control (e.v.). Cells were treated either with DMSO or 250 nM Torin-1 for 16 h and subsequently stained for phospholipids (magenta) and neutral lipids (green) using NileRed. Nuclei were stained with Hoechst (blue); scale bars: 20 µm. (**G)** Western blot analysis of re-expressed, HIS-tagged APLA1 (APLA1 re) and the inactive S111A variant (S111A re). GAPDH was used as a loading control. (**H**) Quantitative analysis of phospholipid deposits and lipid droplets from images shown in (F) (n = 3 fields, data representative of two independent experiments). (**I**) Electron micrographs of HEK293 WT, *APLA1*-KO, *PLA2G15*-KO and dKO cells treated with 250 nM Torin-1 for 16 h; scale bars: 2 µm (overview), 0.5 µm (zoom). (**J**) Analysis of multilamellar body (MLB) diameter from micrographs in (D) (n = number of lysosomes measured). Statistically significant differences were determined by multiple *t*-tests (A), two-way ANOVA with Sidák post-hoc test (D, E, H), and one-way ANOVA with Dunett’s post hoc analysis (D, J, *: compared to DMSO, ^#^: compared to WT) (\**p*<0.05, **^,##^*p*<0.01, \*\*\**p*<0.001).

Lysosomal metabolism is regulated via the mammalian target of rapamycin complexes 1 and 2 (mTORC1 and mTORC2). Pharmacological inhibition of mTORC by Torin-1 promotes membrane trafficking to lysosomes, GPL catabolism, and FA export^6^. Subsequently, liberated FAs are then used for TAG synthesis and stored in lipid droplets (LDs). Previous observations suggested that this process occurs independently of PLA2G15 and that additional lysosomal GPL-degrading enzymes must exist^6^. Accordingly, we investigated whether Torin-1-induced lysosomal GPL turnover and LD formation depend on APLA1. To this end, we treated WT and mutant cells with either DMSO or Torin-1 for 16 h. Cells were subsequently stained with NileRed, which allows simultaneous visualization of phospholipid-rich structures and neutral lipid droplets (LDs) based on distinct excitation/emission spectra (**Fig. S4A, B**)^15,16^.

Under basal conditions, phospholipid deposits were increased in dKO cells, but not in single KO cells, whereas LDs were readily detectable (**Fig. 3C**). Following Torin-1 treatment, phospholipid accumulation was modest in WT and *PLA2G15*-KO cells, but became strikingly pronounced in *APLA1*-KO and dKO cells (**Fig. 3C, D**). Conversely, Torin-1 robustly stimulated LD formation in WT and *PLA2G15*-KO cells, whereas this response was blunted in *APLA1*-KO and dKO cells, consistent with impaired GPL catabolism and FA flux into TAGs (**Fig. 3C, E**). Co-staining with LysoSensor confirmed that the accumulated phospholipid deposits localized to acidic compartments (**Fig. S5A, B**). Stable re-expression of active APLA1, but not the catalytically inactive S111A mutant ameliorated Torin-1-induced phospholipidosis in *APLA1*-KO cells (**Fig. 3F, G**) while fully restoring LD formation (**Fig. 3H**).

Ultrastructure analysis revealed that Torin-1 increased the formation of multilamellar bodies in cells of all genotypes (**Fig 3I**). Lysosomes of *APLA1*-KO and dKO cells exhibited an excessive accumulation of multilamellar membranes, a hallmark of phospholipidosis^17^. In line, the diameter of multilamellar bodies observed in *APLA1*-KO and dKO cells increased ∼2-fold compared to WT controls, while those of *PLA2G15*-KO cells were only moderately increased (**Fig 3J**).

To confirm the impaired lysosomal GPL-breakdown and TAG formation, we determined the flux of GPL-derived FAs into LDs of *APLA1*-KO and dKO cells using fluorescent or radiolabeled FAs as tracers. To label lipid pools, cells were incubated with tracers for 5 h, followed by a 16 h chase period in the presence of Torin-1 (**Fig. 4A**). Under the applied conditions, BODIPY-palmitate (BODIPY FL C16) stained LDs of WT and *PLA2G15*-KO cells, but remained trapped in phospholipid deposits of *APLA1*-KO and dKO cells (**Fig. 4B, C**). Using ^3^H-labeled oleic acid as tracer, we found that the incorporation into total cellular lipids was unchanged between genotypes and slightly increased in response to Torin-1 treatment (**Fig. 4D**). However, Torin-1 promoted ^3^H-TAG synthesis in WT and *PLA2G15*-KO, but not in *APLA1*-KO and dKO cells (**Fig. 4E**).

**Figure 4:**
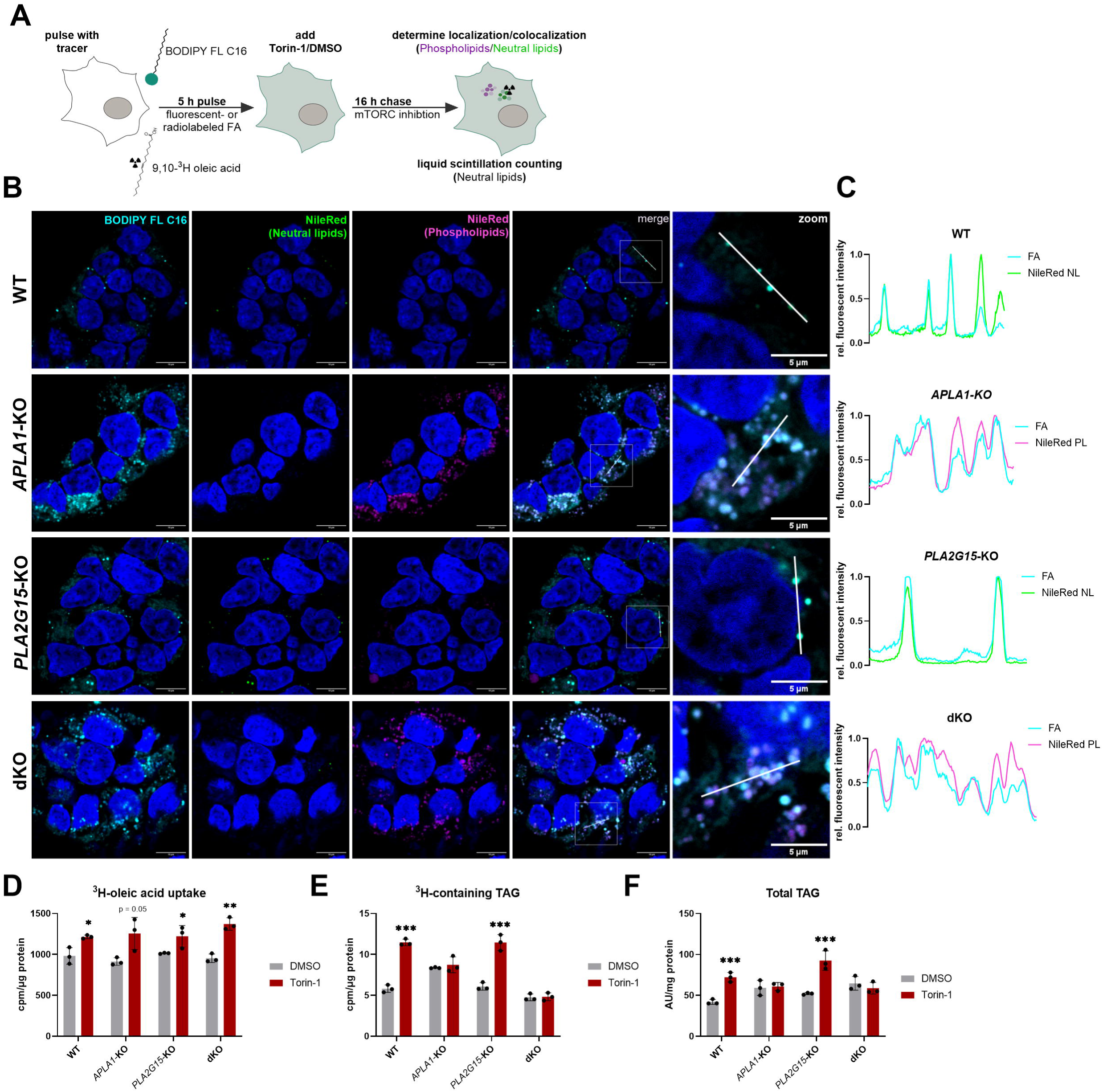
*APLA1*-deficient cells display an impaired flux of phospholipid-derived FAs into TAGs. (**A**) Schematic workflow for pulse-chase experiments using a BODIPY-labeled fatty acid (BODIPY FL C16) or 9, 10-^3^H oleic acid as tracers. (**B**) HEK293 WT and mutant cells were pulsed with 4 µM BODIPY-palmitate (cyan) for 5 h. Cells were then washed and the medium was replaced with standard growth medium containing either DMSO or 250 nM Torin-1. Subsequently, cells were fixed and stained with NileRed (phospholipids: magenta; neutral lipids: green) and Hoechst (blue) for confocal microscopy; scale bars: 10 µm (overview), 5 µm (zoom) (n = 3 fields, data representative of two independent experiments). (**C**) Co-localization of BODIPY FL C16 and NileRed-stained organelles shown in (B) was confirmed by intensity plots. (**D, E**) HEK293 WT and mutant cells were pulsed with 25 nM ^3^H-labeled oleic acid as described in (B). The incorporation of radioactivity into total lipids was determined by liquid scintillation counting. Total triacylglycerols (TAGs) were separated from other lipids by thin layer chromatography. The TAG band was cut out and the co-migrating radioactivity was determined by liquid scintillation counting (n = 3 independent replicates). (**F**) TAG levels in HEK293 WT and mutant cells treated with DMSO or 250 nM Torin-1 for 16 h. TAGs were analyzed by targeted LC-MS (n = 3, data representative of three independent experiments). Statistically significant differences were determined by two-way ANOVA and corrected for multiple comparisons by Sidák’s post-hoc test (\**p*<0.05, \*\**p*<0.01, \*\*\**p*<0.001).

Finally, we determined the cellular TAG content by LC-MS. Torin-1 treatment increased the total TAG content of WT and *PLA2G15*-KO cells, but not of *APLA1*-KO and dKO cells (**Fig. 4F**). Analysis of TAG composition by mass spectrometry revealed an increase in the most abundant TAG species in WT and *PLA2G15*-KO cells (**Fig. S6A**). Thus, the effects of Torin-1 on lysosomal FA mobilization and TAG formation in HEK293 cells depend on APLA1.

In addition to HEK293 cells, we observed Torin-1-induced phospholipidosis in HAP1 (**Fig. S7A, C, D**) and HeLa cells lacking APLA1 (**Fig. S7B, F, G**). In contrast to HEK293 and HAP1 cells, however, Torin-1 treatment did not increase LD formation in HeLa cells (**Fig. S7H**). *APLA1*-KO HeLa cells displayed elevated TAGs under basal conditions, which decreased in response to Torin-1 treatment, indicating differences in FA metabolism between the investigated cell lines. Overall, however, our observations suggest a pivotal role of APLA1 in lysosomal GPL catabolism.

### APLA1 and PLA2G15 are the major lysosomal glycerophospholipases

To study the relative contributions of phospholipases to acid GPL hydrolysis, we first determined acid phospholipase activity in whole cell lysates of WT and mutant cells (**Fig. 5A**). Hydrolysis of PG, PI, and PC in lysates of *APLA1*-KO cells was reduced by 40%, 70%, and 60%, respectively, compared to WT controls. Lysates of *PLA2G15*-KO cells displayed reduced PG, PI, and PC hydrolysis by 40%, 40%, and 20%, respectively. dKO lysates exhibited severely reduced activity against PG and PC (> 90%), while residual activity of ∼30% was detected for PI (**Fig. 5A**). Reduced acid GPL hydrolase activity was also detected in lysates of HAP1 and HeLa cells lacking APLA1 (**Fig. S8A, B**).

**Figure 5:**
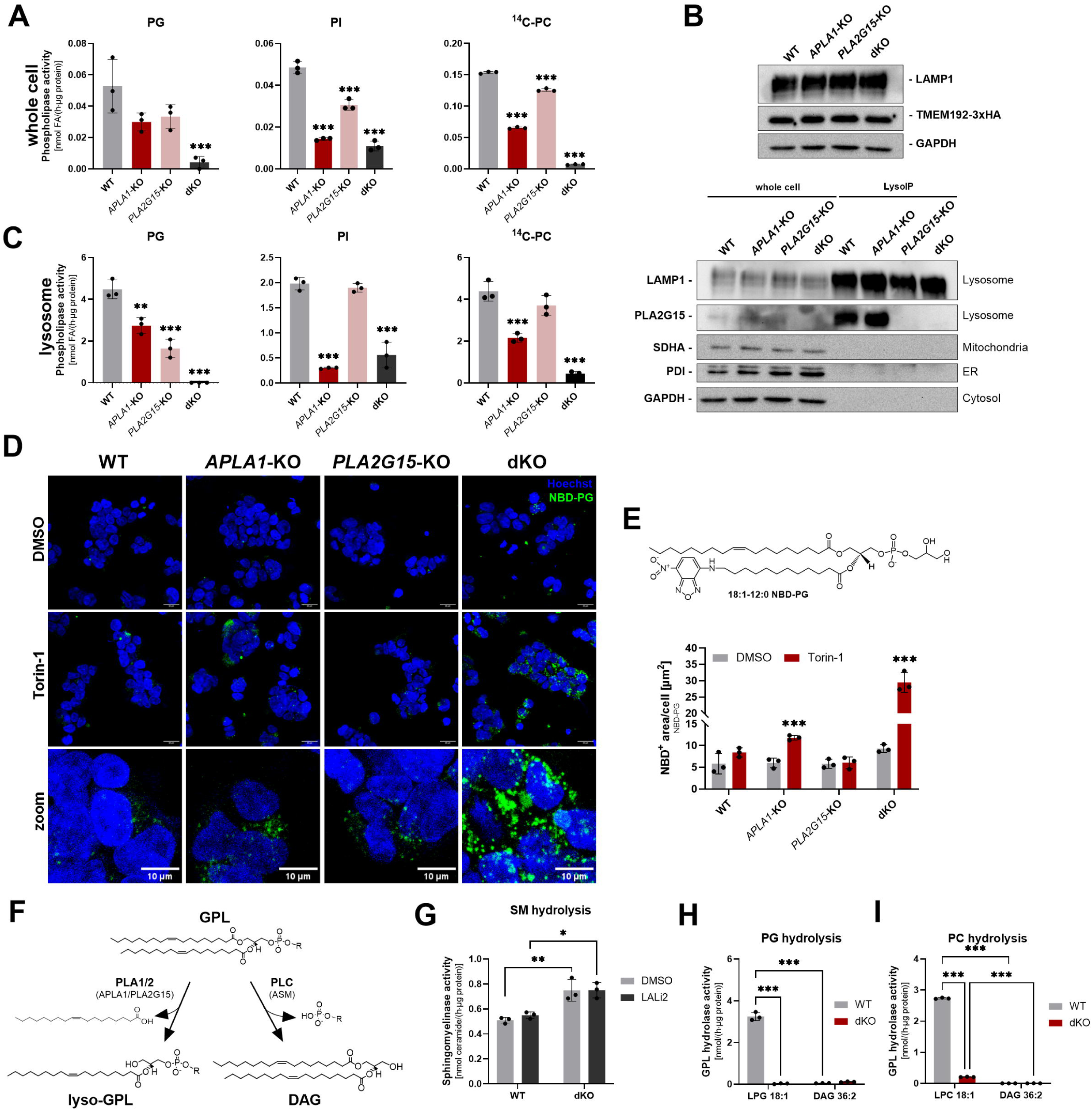
APLA1 and PLA2G15 are the major acid phospholipases A1/2. (**A**) Acid phospholipase activity toward different GPL substrates in whole cell lysates. Lysates of whole cell fractions (0.67 mg/ml) were incubated with 0.33 mM of either PG, liver PI, or a mixture of PC:BMP (70/30 mol%) at pH 4.5 for 1 h. Hydrolysis of PG and PI was assessed by measuring liberated fatty acids using an enzymatic kit. For PC hydrolysis, unlabeled PC (final concentration: 233 µM) was mixed with di-oleoyl-1-^14^C,^14^C-PC, (final concentration: 6 µM) and unlabeled BMP (30 mol%). The release of ^14^C oleic acid was quantified by liquid scintillation counting (n = 3 independent replicates, data representative of two independent experiments with at least two knock-out clones per genotype). (**B**) Lysosomes were isolated from cells expressing 3xHA-tagged TMEM192 (top) by LysoIP. Purity was assessed by Western blot analysis of equal volumes using LAMP1 (lysosome, membrane), PLA2G15 (lysosome, lumen), SDHA (mitochondria), PDI (ER), and GAPDH (cytosol) as marker proteins (bottom). (**C**) Acid phospholipase activity assays in lysosomal preparations. Assays were performed as described in (A) using 0.007 mg/ml protein for PG and PC hydrolysis and 0.017 mg/ml protein for PI hydrolysis) (n = 3 independent replicates, data representative of two independent experiments). (**D**) Cells were labeled with 2 µM NBD-PG (green) for 4 h. Subsequently, the medium was replaced with medium containing either DMSO or 250 nM Torin-1 for 16h. Cells were then fixed and analyzed by confocal microscopy. Nuclei were stained with Hoechst (blue); scale bars: 20 µm (overview), 10 µm (zoom). (**E**) Quantitative analysis of NBD-positive (NBD^+^) area from images shown in (D) (n = 3 fields). (**F**) Schematic illustration of lysosomal GPL-degradation pathways mediated by PLA1/2 and PLC, yielding lyso-GPLs and diacylglycerols (DAGs), respectively. (**G**) ASM (PLC) activity in lysosomal lysates of HEK293 WT and dKO cells. Activity was determined as ceramide formation from sphingomyelin (isolated from porcine brain). Lysosomal lysates (0.01 µg/µl) were incubated with 0.4 mM SM at pH 5 and 37°C for 1 h and the formation of ceramides was determined by targeted MS-analysis (n = 3 independent replicates). Lysosomal lysates were preincubated with the LIPA-inhibitor LAListat2 (LALi2, 1 µM) for 10 min at room temperature. (**H** and **I**) Breakdown of PG and PC into lyso-GPLs and DAG by lysosomal phospholipases. Lysosomal lysates (0.01 mg/ml) of HEK293 WT and dKO cells were incubated with 400 µM PG or a mixture of PC/BMP (70/30 mol%) under the conditions described in (G). The formation of lyso-GPLs and DAG was analyzed by LC-MS (n = 3 independent replicates). Statistically significant differences were determined by one-way ANOVA corrected for multiple comparisons by Dunnett’s post-hoc analysis (A and C) and by two-way ANOVA corrected for multiple comparisons by Tukey’s post-hoc test (E, G, H, I) (\**p*<0.05, \*\**p*<0.01, \*\*\**p*<0.001).

Next, we monitored GPL hydrolysis in purified lysosomes of HEK293 cells. For this purpose, we stably expressed the HA-tagged integral lysosomal membrane protein TMEM192 in cells and purified organelles by lysosome immunopurification (LysoIP)^18^, yielding a strong enrichment of lysosomes with minimal impurities from other compartments (**Fig. 5B**). This procedure led to an ∼100-fold enrichment in acid PG hydrolase activity relative to whole cell fractions (**Fig. 5A** versus **5C**). Consistent with whole-cell measurements, lysosomal lysates from mutant cells showed markedly reduced phospholipase activity toward PG, PI, and PC, compared to WT controls (**Fig. 5C**).

Notably, PG hydrolase activity was virtually not detectable in lysosomal lysates of dKO cells. Based on this observation, we treated cells with fluorescently labeled phosphatidylglycerol (NBD-PG) to assess lysosomal PG accumulation. No substantial accumulation was observed under basal conditions (DMSO, **Fig. 5D, E**). However, in accordance with the almost complete lack of acid PG hydrolase activity, stimulation of lysosomal phospholipid turnover with Torin-1 resulted in a strong accumulation of the tracer in dKO cells (**Fig. 5D, E**). Torin-1 also induced a moderate increase in *APLA1*-KO, but not *PLA2G15*-KO cells, suggesting that the presence of APLA1 is sufficient to prevent lysosomal PG accumulation.

In addition to PLA1/2-dependent mechanisms, GPL degradation may also be initiated by phospholipase C (PLC), leading to the formation of DAG and phosphorylated head groups (**Fig. 5F**). Monitoring ceramide formation from sphingomyelin clearly revealed PLC activity in lysosomal preparations, catalyzed by acid sphingomyelinase (ASM), the only known lysosomal PLC^19^. ASM activity was increased by ∼30% in dKO samples in comparison to the WT control (**Fig. 5G**). To investigate whether ASM contributes to GPL degradation, we compared PLA1/2 and PLC activity by measuring lyso-GPL and DAG formation. To avoid further hydrolysis of DAG by lysosomal acid lipase (LAL), we conducted reactions in the presence of the LAL inhibitor Lalistat-2 (LALi2)^20^, which did not affect ASM activity (**Fig. 5G**). Using PG or PC/BMP as substrate, PLA1/2-mediated formation of LPG and LPC was readily detected in lysosomal preparations of WT cells, but was severely reduced in lysates of dKO cells (**Fig. 5H, I**). In contrast, DAG generation from PG or PC was minimal, regardless of the genotype. These observations suggest that PLC does not substantially contribute to lysosomal GPL degradation, even in cells lacking acid PLA1/2 activity.

### Loss of APLA1 results in the accumulation of multiple GPL species within lysosomes

To investigate the impact of *APLA1*# and *PLA2G15*-deficiency on lysosomal lipid composition, we performed untargeted LC–MS analysis of purified lysosomes from WT, *APLA1*-KO, *PLA2G15*-KO, and dKO cells under basal conditions and following Torin-1 treatment. Principal component analysis revealed that Torin-1 induced a remodeling of the lysosomal lipidome across all genotypes, consistent with enhanced lysosomal lipid turnover (**Fig. 6A**, top). Despite this common response, *PLA2G15*-KO lysosomes clustered closely with WT lysosomes under both conditions, whereas dKO lysosomes formed a distinct cluster, indicative of a profound disruption of lysosomal GPL metabolism. *APLA1*# KO lysosomes remained similar to WT lysosomes under basal conditions but separated following Torin-1 treatment (**Fig. 6A**, top). Loading plot analysis revealed that the observed separation between the genotypes was primarily driven by alterations in GPL species (**Fig. 6A**, bottom).

**Figure 6:**
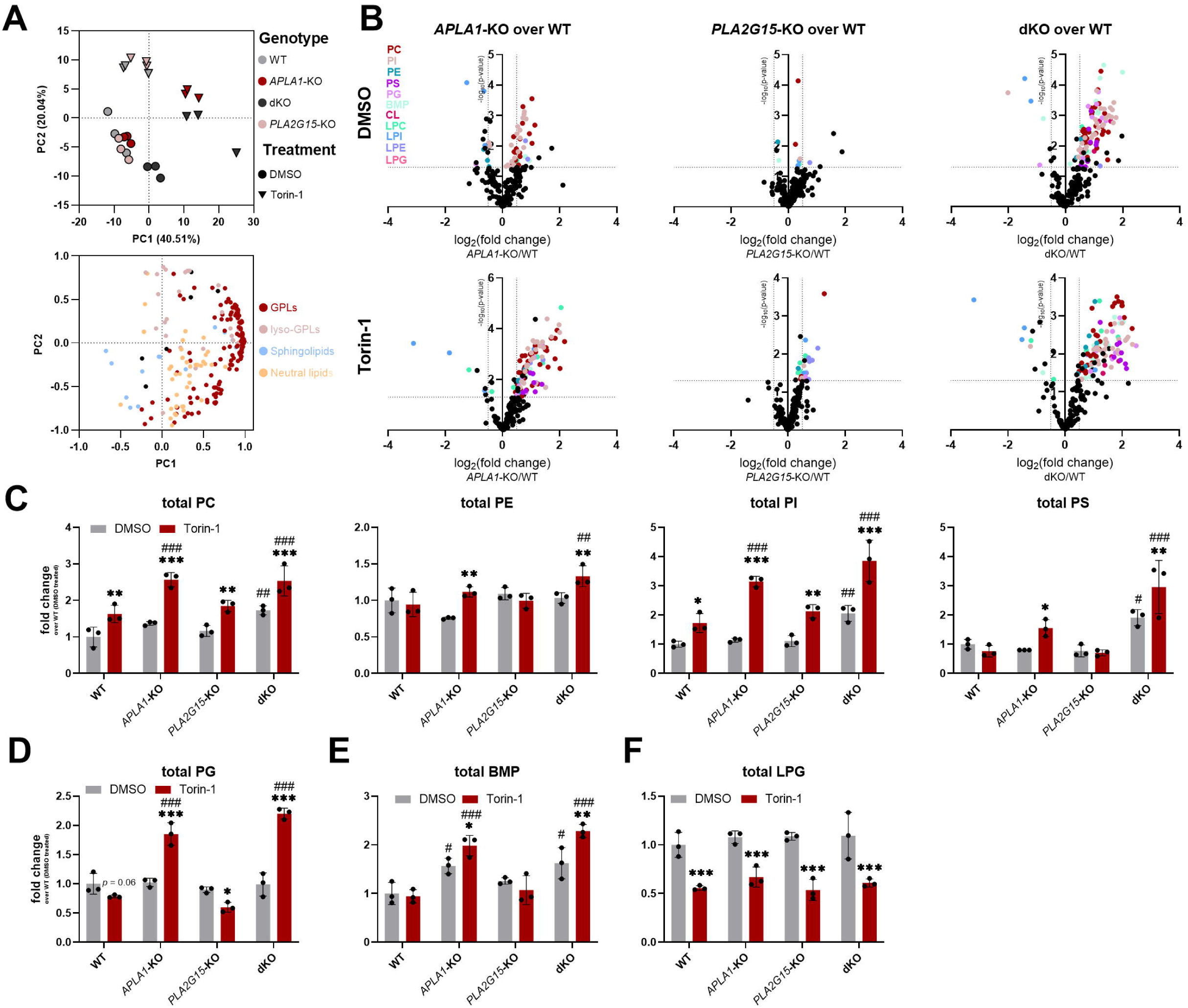
*APLA1*-deficient HEK293 cells accumulate GPLs in lysosomes upon Torin-1 treatment. (**A**) Principal component analysis (upper graph) with corresponding loadings (lower graph) of lipidomic data from isolated lysosomes of HEK293 WT and KO cells treated with DMSO or 250 nM Torin-1 (n = 3 biological replicates, data representative of two independent experiments). (**B**) Volcano plots of lipidomic data from (A). Data are represented as log_2_-fold changes over WT lysosomes treated with DMSO (upper row) or Torin-1 (lower row) (n = 3 biological replicates). (**C**) Total PC, PE, PI, and PS levels in lysosomes of WT and mutant cells based on data from (A). (**D-F**) Targeted analysis of PG (D), BMP (E), and LPG (F) levels in lysosomes isolated from WT and mutant cells treated with either DMSO or Torin-1 for 16 h (n = 3 biological replicates). Data are represented as mean (A, B) and mean ± SD (C, D, F). Statistically significant differences were determined by Student’s unpaired two-tailed *t*-tests (B) and two-way ANOVA corrected for multiple comparisons by Sidák’s post-hoc test (C, D, E, F) (*: DMSO vs. Torin-1, ^#^: compared to WT, *^,#^*p*<0.05, **^,##^*p*<0.01, ***^,###^*p*<0.001).

Consistent with the PCA analysis, direct investigation of lysosomal lipid species revealed a progressive accumulation of GPLs upon loss of APLA1. Under basal conditions, *APLA1*-KO and dKO lysosomes exhibited increased abundance of 28% and 70% of detected GPL species, respectively, compared with WT lysosomes (**Fig. 6B**). Following Torin-1 treatment, GPL accumulation was further exacerbated in *APLA1*-KO and dKO lysosomes, whereby 70% of the detected GPL species increased in both genotypes (**Fig. 6B**). All major GPL classes showed significant changes (**Fig. 6C**). The most prominently affected lipid classes were phosphatidylcholine (PC) and phosphatidylinositol (PI), with alterations observed across the majority of detected species (**Fig. 6C**; **Tables 1–3**). In contrast, *PLA2G15*-KO lysosomes displayed only modest changes in GPL composition compared to WT lysosomes under both basal and Torin-1 conditions, consistent with previous observations and further supporting the central role of APLA1 in lysosomal GPL turnover.

Previous studies proposed that lysosomal BMP synthesis depends on acid PG hydrolases generating LPG, the substrate of the lysosomal BMP synthase ceroid lipofuscinosis neuronal 5 (CLN5)^21,22^. To test whether loss of PG hydrolase activity affects PG, LPG, and BMP levels, we employed targeted LC-MS on isolated lysosomes to accurately discriminate between PG and BMP isomers. PG levels were unchanged under basal conditions and increased upon Torin-1 treatment in *APLA1*-KO and dKO lysosomes, in line with impaired hydrolysis (**Fig. 6D**). BMP levels were elevated in *APLA1*-KO and dKO samples in the absence and presence of Torin-1 (**Fig. 6E**). However, despite the severe reduction in acid PG hydrolase activity in dKO cells, lysosomal LPG levels were similar across all genotypes (**Fig. 6F**), indicating that BMP synthesis does not require lysosomal PG hydrolysis. These findings are consistent with recent reports suggesting that the BMP precursor LPG is produced at the ER by the Batten disease protein ceroid lipofuscinosis neuronal 8 (CLN8)^23,24^.

### APLA1 is inhibited by cationic amphiphilic drugs

Cationic amphiphilic drugs (CADs) cause lysosomal phospholipid accumulation, a pathological condition called drug-induced phospholipidosis (DIPL)^17^. To investigate whether APLA1 is sensitive to CADs, we first tested the effects of amiodarone and chloroquine on acid phospholipases, as these substances are prototypical DIPL inducers. The addition of amiodarone to lysosomal extracts from WT cells completely inhibited acid phospholipase activity against PC/BMP (70/30 mol%), similar to what was observed in dKO samples, suggesting inhibition of both APLA1 and PLA2G15 activity (**Fig. 7A**). Using PC/BMP or PE/BMP (70/30 mol%) as substrates, we determined IC_50_ values for amiodarone-mediated APLA1 inhibition of 44 µM and 36 µM, respectively (**Fig. 7B**). In comparison to amiodarone, chloroquine was less potent with IC_50_ values of 293 and 96 µM, respectively (**Fig 7D**). When PG was used as a substrate, amiodarone inhibited APLA1 activity with an IC value of 112.5 µM, indicating that substrate charge correlates CAD efficacy (**Fig. 7B**). Consistent with a charge-based mechanism, liposome pulldown experiments revealed that amiodarone blocked APLA1 binding to PC/BMP liposomes (**Fig 7C**, Inset).

**Figure 7:**
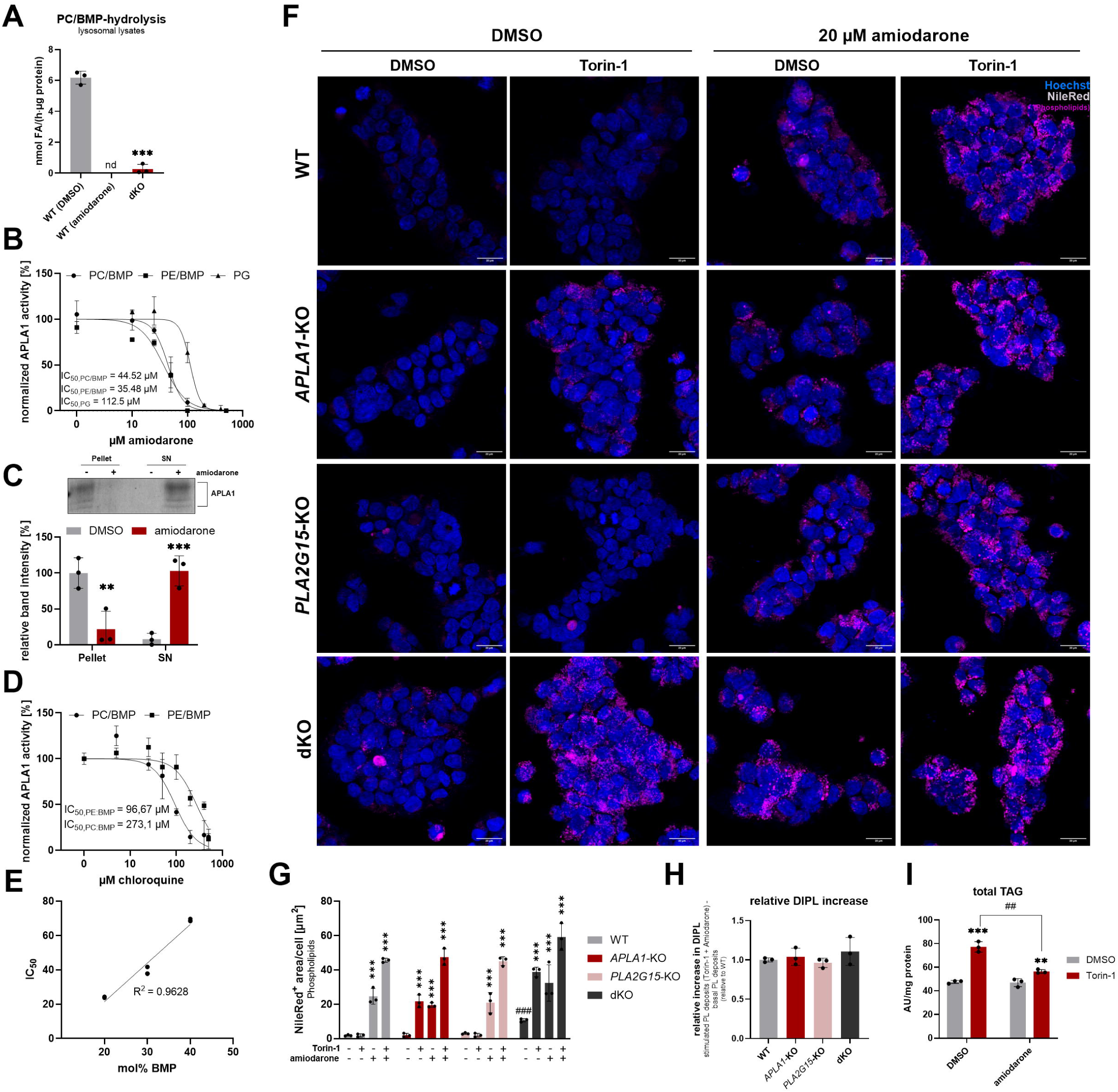
APLA1 and PLA2G15 are inhibited by cationic amphiphilic drugs. Phospholipase activity was measured as the release of FAs using a colorimetric assay with 0.4 mM phospholipid substrates at pH 4.5 for 1h. (**A**) Lysosomal lysates from HEK293 WT and dKO cells (0.008 mg/ml protein) were incubated with 0.4 mM PC/BMP (70/30 mol%) in the absence (DMSO) or presence of 50 µM amiodarone. (n = 3 independent replicates, data representative of two independent experiments). (**B**) Dose-dependent inhibition of purified APLA1 by amiodarone. APLA1 (0.3 µM) was incubated with 0.4 mM PC/BMP, PE/BMP (70/30 mol%), or PG in the presence of the indicated concentrations of amiodarone (n = 3 independent replicates). (**C**) Amiodarone prevents APLA1 binding to its substrate. Purified APLA1 (3 µM) was incubated with 2 mM PC/BMP (70/30 mol%) at pH 4.5 in the presence or absence of 300 µM amiodarone. Samples were centrifuged at 100,000 x g. Subsequently, pellet and supernatant (SN) fractions were analyzed by SDS-PAGE and Coomassie staining (data are representative of three independent experiments). (**D**) Dose-dependent inhibition of purified APLA1 by chloroquine. APLA1 (0.3 µM) was incubated with 0.4 mM PC/BMP or PE/BMP (70/30 mol%) in the presence of the indicated concentrations of chloroquine (n = 3 independent replicates). (**E**) Amiodarone-mediated inhibition of APLA1 depends on substrate charge. Purified APLA1 (0.3 µM) was incubated with PC/BMP substrate containing the indicated mol% of BMP in the presence of increasing concentrations of amiodarone. IC_50_ values were determined as shown in (B) and plotted against BMP concentrations (n = 3 independent replicates, data representative of two independent experiments). (**F**) WT and mutant HEK293 cells develop phospholipidosis to a similar extent following treatment with amiodarone and Torin-1. Cells were treated with DMSO, 20 µM amiodarone, 250 nM Torin-1 or a combination thereof and phospholipid deposits (magenta) were stained with NileRed. Nuclei were stained with Hoechst (blue); scale bars: 20 µm. (**G**) Quantitative analysis of NileRed-positive phospholipid deposits from (F) (n = 3 fields). (**H**) Relative increase in drug-induced phospholipidosis in WT and mutant cells in response to combined treatment with amiodarone and Torin-1 (calculated from data shown in (G)). (**H**) Amiodarone prevents Torin-1-induced TAG accumulation in WT cells. HEK293 cells were cultured with 20 µM amiodarone or DMSO in the absence and presence of 250 nM Torin-1 for 16 h. Total TAG levels were determined by targeted MS-analysis (n = 3 independent replicates). Statistically significant differences were determined by Student’s unpaired two-tailed *t*-tests (A), one-way ANOVA corrected for multiple comparison by Dunnett’s post-hoc test (G: ^#^) and two-way ANOVA corrected for multiple comparisons by Tukey’s (G: *) and Sidák’s (C, I) post-hoc tests (**^,##^*p*<0.01, ***^,###^*p*<0.001).

Notably, IC□□ values determined for PC/BMP substrates positively correlate with their BMP content (**Fig. 7E**). These results support the concept that amiodarone masks the negative charge of the substrate, which is crucial for the activation of APLA1, rather than directly inhibiting the enzyme’s catalytic function.

To monitor DIPL, we treated HEK293 cells with amiodarone in the absence and presence of Torin-1 and visualized phospholipid deposits with NileRed (**Fig. 7F**). Amiodarone, but not Torin-1, increased phospholipidosis in WT cells, while combined treatment doubled phospholipid accumulation (**Fig. 7G**). *PLA2G15*-KO cells behaved similarly as WT cells (**Fig. 7G**). In *APLA1*-KO cells, both amiodarone and Torin-1 treatment increased phospholipidosis. Combined treatment in *APLA1*-deficient cells promoted phospholipid accumulation to levels of double-treated WT cells (**Fig. 7G**). dKO cells showed phospholipidosis under basal conditions, which further increased in response to amiodarone or Torin-1 treatment. Combined treatment of dKO cells had again an additive effect, resulting in moderately increased phospholipid accumulation compared to WT cells treated with both compounds (**Fig. 7G**). However, the relative increase over basal levels upon combined treatment was comparable across all genotypes, suggesting that amiodarone treatment of WT cells causes a phospholipidosis phenotype similar to that of dKO cells (**Fig. 7H**). Finally, amiodarone blunted the Torin-1-induced increase in TAG formation in WT cells, a process dependent on APLA1 (**Fig. 7I**). Overall, these findings suggest that CAD-mediated inhibition of both APLA1 and PLA2G15 contributes to the development of DIPL.

## Discussion

PPT2/APLA1 was originally identified as a homolog of PPT1, which cleaves thioester bonds of S-palmitoylated proteins in lysosomes^12^. Mutations in *PPT1* cause an early-onset form of Batten disease, ceroid lipofuscinosis neuronal 1 (CLN1), which is characterized by the accumulation of ceroid-lipofuscin pigments in lysosomes of neurons and other cell types^25^. Although, PPT1 and PPT2/APLA1 were initially thought to exhibit similar activities, subsequent structural and biochemical work demonstrated that APLA1 cannot deacylate proteins due to steric restrictions in its active site, leaving its cellular function elusive^12,13^. Cases of *APLA1* deficiency in humans have not been documented. Importantly, however, *Ppt2*/*Apla1*-deficient mice develop a late-onset Batten disease-like phenotype, characterized by autofluorescent bodies throughout the brain, progressive motor abnormalities, and neurodegeneration^26,27^. In addition, massive storage of ceroid-lipofuscin pigments has been observed in peripheral tissues, including the spleen, exocrine pancreas, and bone marrow^27^, suggesting that APLA1 is broadly expressed and essential for proper lysosomal function.

Here, we demonstrate that APLA1 degrades GPLs in concert with PLA2G15. *APLA1*-deficiency impairs lysosomal membrane clearance in human cells, despite the overlapping substrate specificity of APLA1 and PLA2G15. Under standard cell culture conditions, phospholipidosis was barely detectable in *APLA1*-KO cells, likely due to low lysosomal membrane catabolism. However, stimulation of GPL turnover by pharmacological inhibition of mTORC led to massive accumulation of GPLs in lysosomes and reduced LD formation, consistent with a requirement for APLA1 in GPL hydrolysis and FA supply for TAG synthesis. In accordance with a previous study, we found that the Torin-1-induced effects on lysosomal GPL degradation and TAG storage are independent of PLA2G15^6^. However, combined deletion of both enzymes causes a severe reduction in acid phospholipase activity and exacerbates phospholipidosis, suggesting that APLA1 and PLA2G15 are the major lysosomal PLA1/2. Our data further indicate that PLC plays only a minor role in lysosomal GPL catabolism, although ASM # the only known acid PLC # is capable of hydrolyzing GPLs in vitro^9^.

Lumenal membranes of late endosomes and lysosomes are enriched in BMP, promoting the activity of lysosomal hydrolases and lipid transfer proteins^14,28,29^. We show that APLA1 requires electrostatic interactions for efficient hydrolysis of GPLs. Unlike anionic GPLs, the hydrolysis of zwitterionic GPLs requires negatively charged BMP as a co-factor to enable APLA1 binding to the substrate. This charge-dependency renders APLA1 sensitive to CADs, which are trapped in lysosomes and interfere with negatively charged intralumenal membranes, ultimately leading to DIPL^17^. CADs comprise a broad spectrum of clinically relevant drugs^30^ and, consequently, avoiding DIPL has become a central objective in drug design. We found that the prototypical CADs amiodarone and chloroquine inhibit APLA1 activity at concentrations corresponding to those previously described for PLA2G15^31^, suggesting that both APLA1 and PLA2G15 are CAD targets.

Both, APLA1 and PLA2G15 can degrade multiple GPL species, including BMP^32,33^. Notably, genetic deletion of PLA2G15 increases BMP levels in mouse tissues and prolongs the lifespan of Niemann-Pick type C1 mice by reducing lysosomal cholesterol burden^7^. Similar beneficial effects were observed after increasing the BMP content in these mice with pharmacological approaches^34^. PLA2G15 degrades BMP containing fatty acids in *sn-3* position (3,3’-*S,S*-BMP and 2,3’-*S,S*-BMP), while the major species 2,2’-*S,S*-BMP is highly resistant to lysosomal hydrolases^7,35^. We found that APLA1 binds to BMP and degrades 3,3’-*S,S*-BMP, but with lower specific activity compared to other anionic substrates. Using mixed substrates (PC/BMP and PE/BMP), APLA1 almost exclusively hydrolyzed zwitterionic GPLs. These observations indicate that, under physiological conditions, BMP primarily serves to recruit and activate APLA1 on intralumenal membranes rather than acting as its main substrate.

BMP synthesis depends on the availability of LPG, which is converted into BMP by CLN5 via a transacylation reaction^22^. Despite the severe reduction in PG hydrolase activity in dKO cells and the accumulation of PG in lysosomes, we detected unchanged lysosomal LPG and increased BMP levels, indicating that the BMP precursor LPG does not originate from lysosomal PG hydrolysis. Recent studies suggest that LPG is produced by the Batten disease protein CLN8 at the ER, transported to lysosomes, and selectively converted into BMP. *CLN8*-deficiency results in severely reduced lysosomal LPG and BMP levels^23,24^. We therefore conclude that the BMP precursor LPG is predominantly supplied via the CLN8 pathway rather than by hydrolysis of PG in lysosomes.

In summary, our study delineates the mechanism of lysosomal GPL catabolism and identifies APLA1, together with PLA2G15, as the major acid PLA1/2 responsible for membrane GPL turnover. These enzymes share substrate specificity but differ in their regiospecificity. Genetic as well as CAD-mediated inactivation of APLA1 compromises membrane catabolism in lysosomes. Consequently, our findings uncover *APLA1* as a candidate risk gene for inherited and drug-induced lysosomal storage disorders.

## Supplemental figure legends

**Figure S1:**
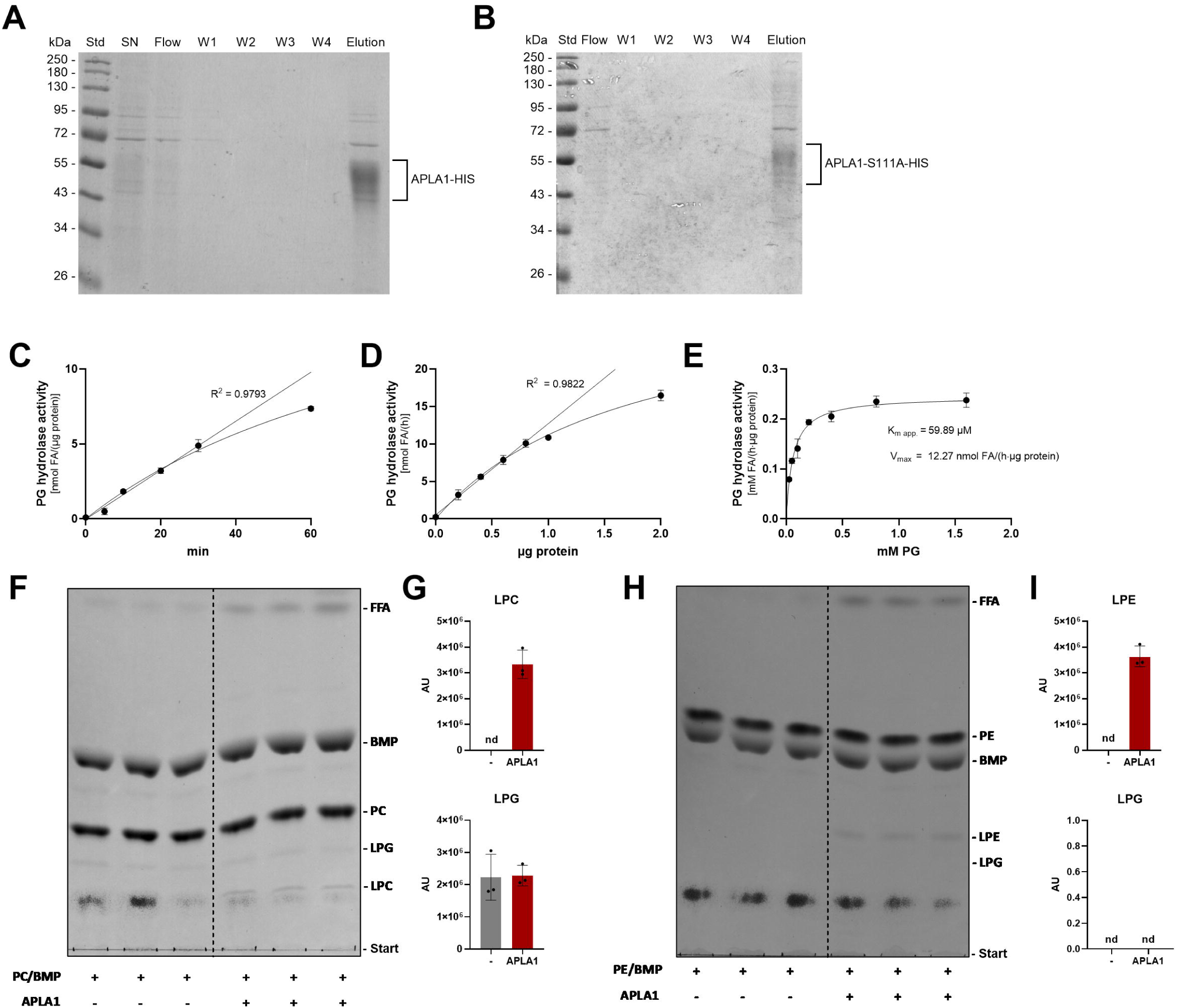
Purification and biochemical characterization of recombinant APLA1 and the inactive S111A mutant. (**A**) HIS-tagged APLA1 and the (**B**) S111A variant were expressed in Expi293F cells and purified from supernatants (SN) by affinity chromatography. Protein fractions were separated by SDS-PAGE and stained with Coomassie blue (flow through (Flow), washing steps (W1-4), eluted protein (Elution)) (**C**) Time-dependent hydrolysis of phosphatidylglycerol (PG) by purified APLA1 at pH 4.5 using 0.5 µg protein and 0.4 mM substrate. (**D**) Dose-dependent hydrolysis of PG by purified APLA1. The enzyme was incubated with 0.4 mM PG for 30 min at pH 4.5. (**E**) Michaelis-Menten kinetics for APLA1-catalyzed PG hydrolysis. Recombinant APLA1 (0.5 µg) was incubated with the indicated substrate concentrations at pH 4.5 for 30 min. In (A)-(E), the release of FAs was determined using an enzymatic kit (n = 3 independent replicates). (**F-I**) Thin layer chromatography of APLA1 reaction products using (F) PC/BMP (70/30 mol%) and (H) PE/BMP (70/30 mol%) as substrates. Bands comigrating with LPC, LPG, and LPE standards were analyzed with Image Lab. Data are presented as mean ± SD (n = 3 independent replicates, data representative of two independent experiments).

**Figure S2:**
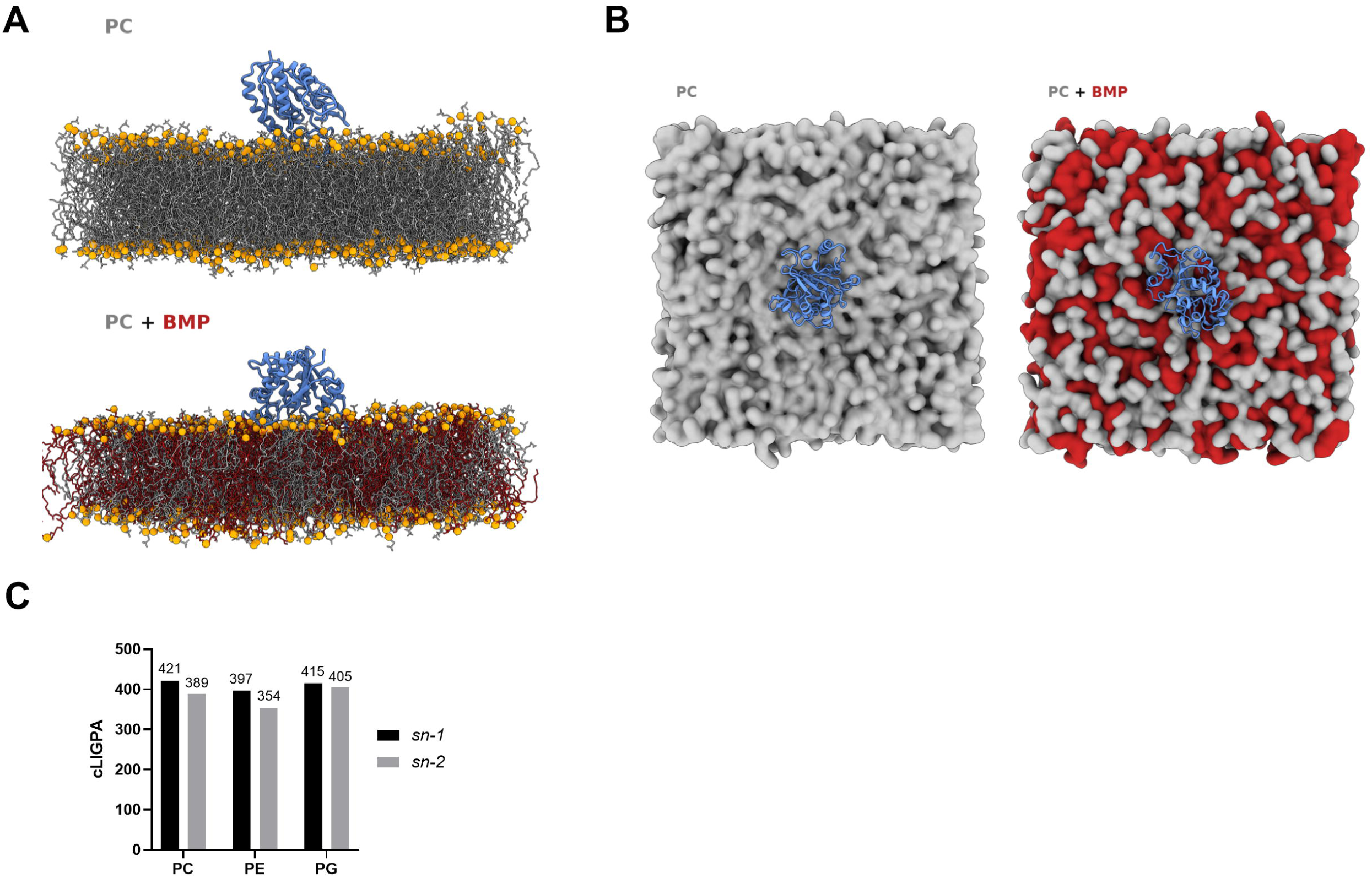
Membrane engagement of APLA1 requires anionic BMP. (**A, B**) Representative snapshots of the two 500 ns equilibrated APLA1-membrane systems (PC versus PC/BMP). Pure PC layer: 768 molecules, PC/BMP layer: 384 PC + 384 BMP molecules. In (A) APLA1 is represented as a blue cartoon, lipid tails are shown as grey (PC) and dark-red sticks (BMP), and phosphate headgroup atoms as orange spheres. Water, ions, and non-polar hydrogens are omitted for clarity. In (B), APLA1 is represented as a blue cartoon and phospholipids are represented as molecular surfaces (gray: PC, dark red: BMP). (**C**) Near attack conformation of PC, PE, and PG predictions using Boltz-2. Bars show confident ligand-protein interface contact pairs (cLIGPA) in the APLA1-PC/PE/PG near-attack complexes involving the active-site serine and the *sn-* 1 and *sn-*2 carbonyl carbon atoms, respectively.

**Figure S3:**
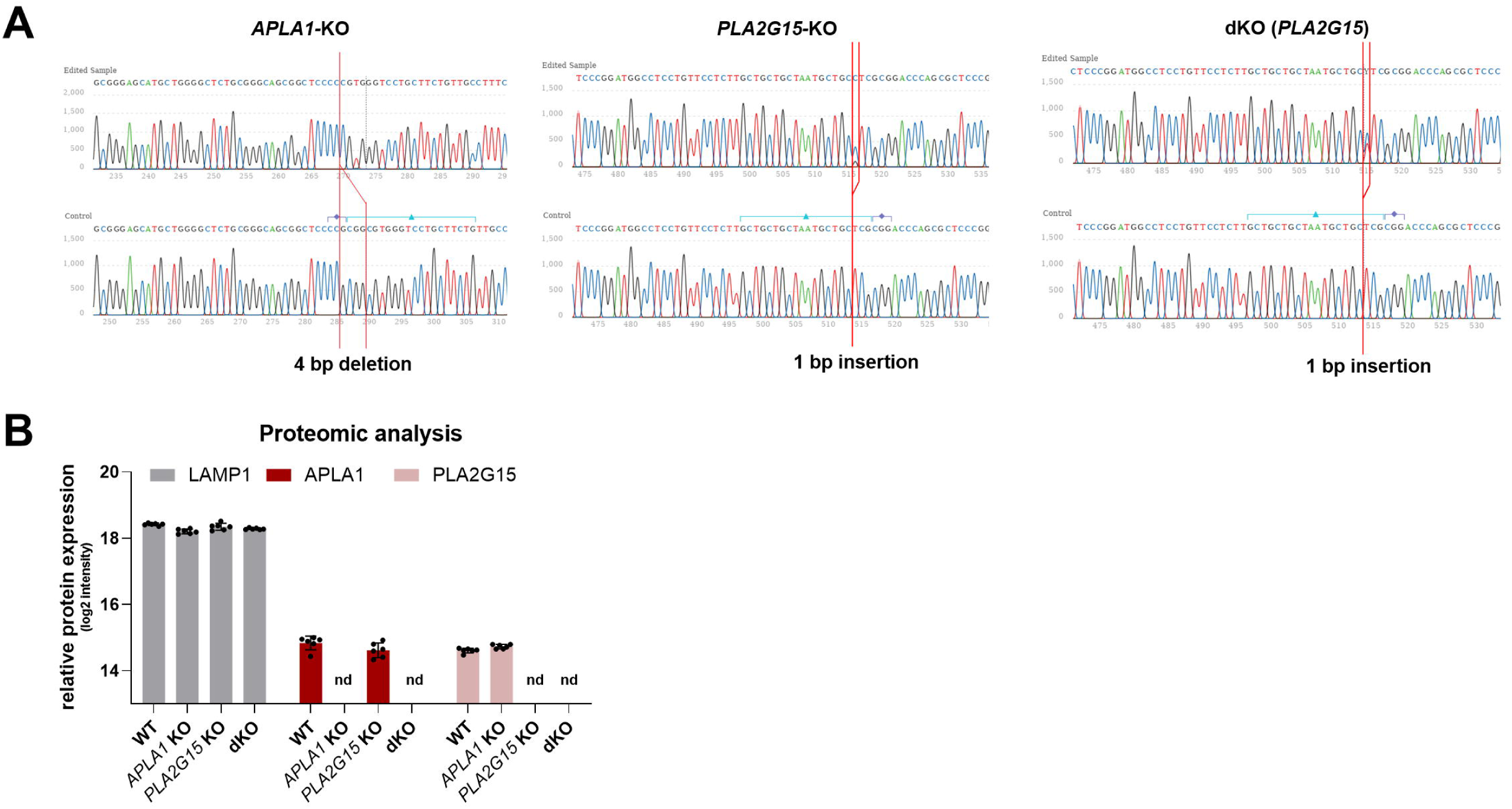
Validation of CRISPR/Cas9-mediated deletion of *APLA1*, *PLA2G15*, or both enzymes (dKO) in HEK293 cells. (**A**) Sanger sequencing and (**B**) proteomic analysis (n = 6 biological replicates, nd: not detected). dKO cells were generated by the transfection of *APLA1*-KO cells with plasmids encoding gRNAs targeting *PLA2G15*.

**Figure S4:**
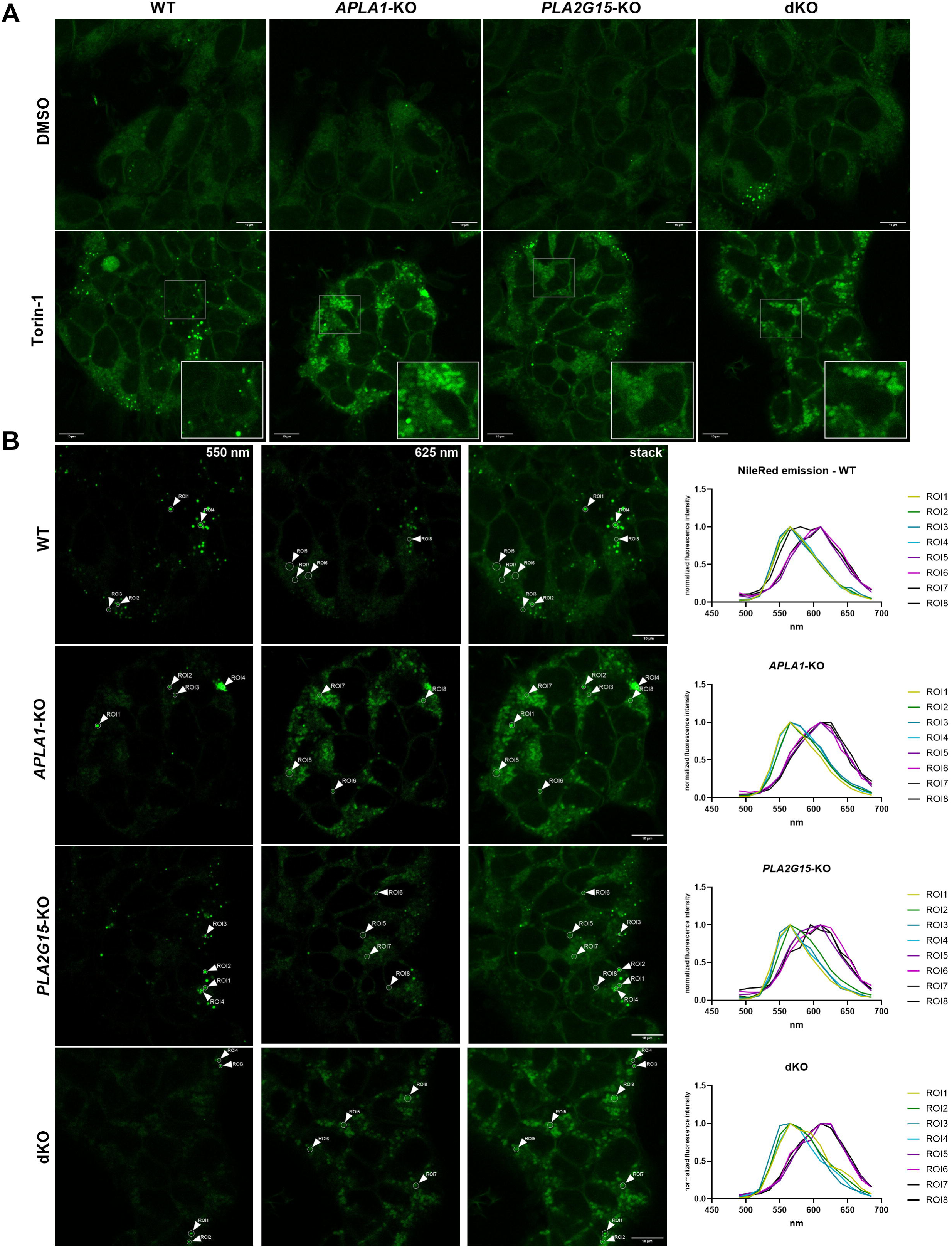
Distinguishing between polar and neutral lipids using Nile Red. **(A)** NileRed emissions (green) of lipid deposits in Torin-1# or DMSO-treated WT and mutant cells. Excitation of NileRed was performed at 488 nm and emission was monitored from 490 to 685 nm. Scale bars: 10 µm. (**B**) Lambda-scan analysis of Torin-1-treated cells. Excitation of NileRed was performed at 488 nm and emission was monitored in 15 nm steps. Images at 550 nm (left panel), 625 nm (middle) and maximum intensity projection of lambda-stacks (right panel) are shown. Arrowheads indicate regions of interest (ROI) for which the wavelength of maximum intensity was determined (right panel). Emission maxima of 550 nm and 625 nm indicate neutral and polar lipid deposits, respectively.

**Figure S5:**
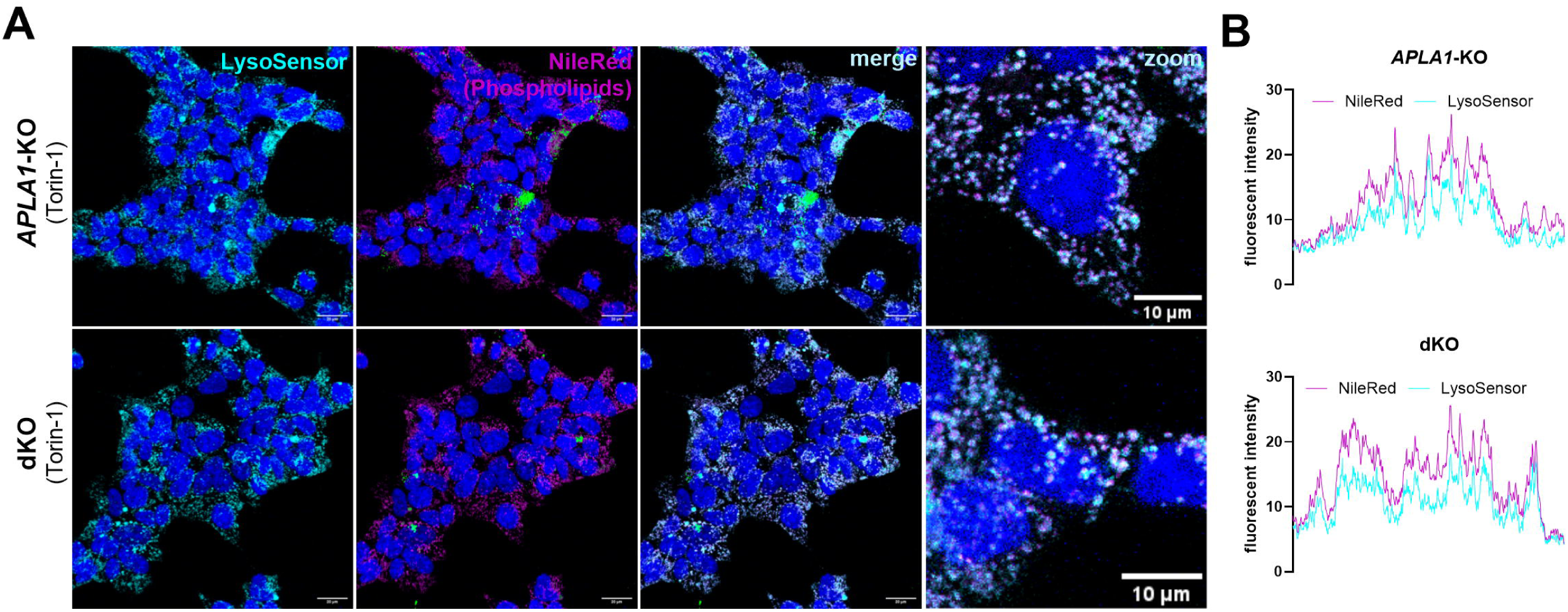
*APLA1*-deficient cells accumulate phospholipids in acidic compartments. (**A**) LysoSensor (cyan) and NileRed (magenta: phospholipids/green: neutral lipids) co-staining of *APLA1*-KO and dKO cells. Nuclei were stained with Hoechst (blue); scale bars: 20 µm (overview), 10 µm (zoom). (**B**) Corresponding intensity plots. Data are representative of two independent experiments.

**Figure S6:**
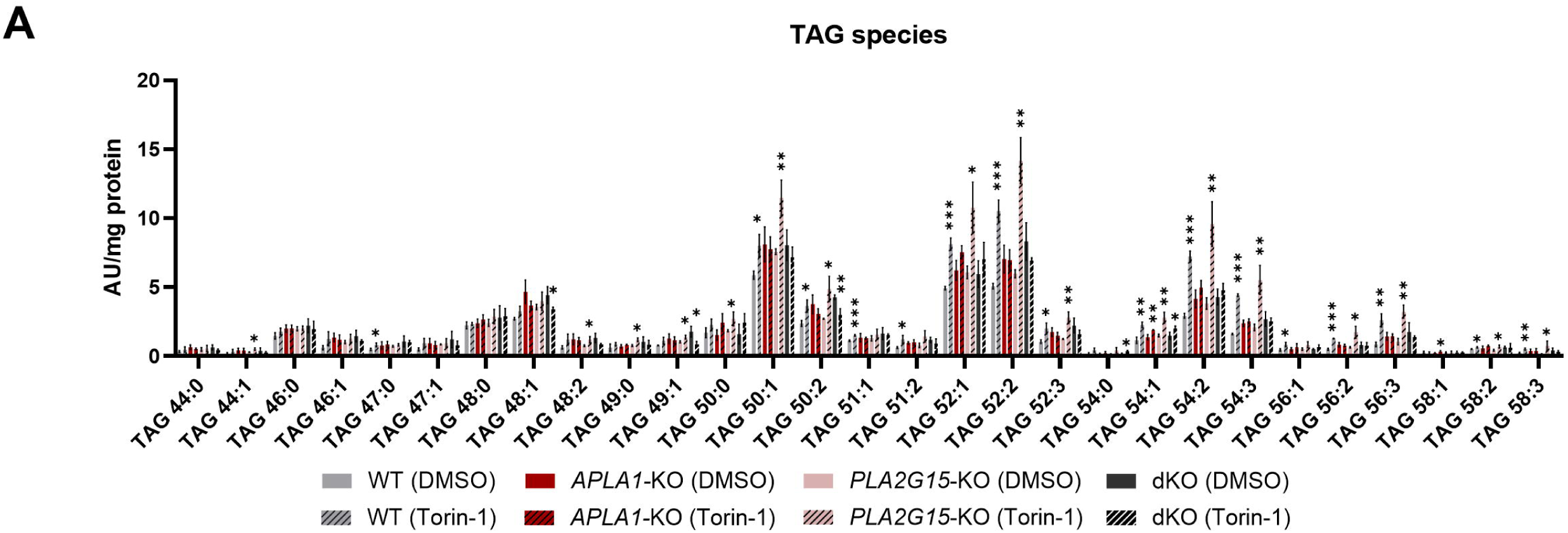
Species-resolved triacylglycerol profiling in HEK293 WT and knockout cell lines following Torin-1 treatment. **(A)** Individual triacylglycerol (TAG) species in HEK293 WT and KO cell lines treated with DMSO (vehicle) or 250 nM Torin-1 for 16 h. TAG levels were determined by targeted LC quadrupole-time of flight MS (LC Q-TOF MS) analysis (n = 3 independent replicates).

**Figure S7:**
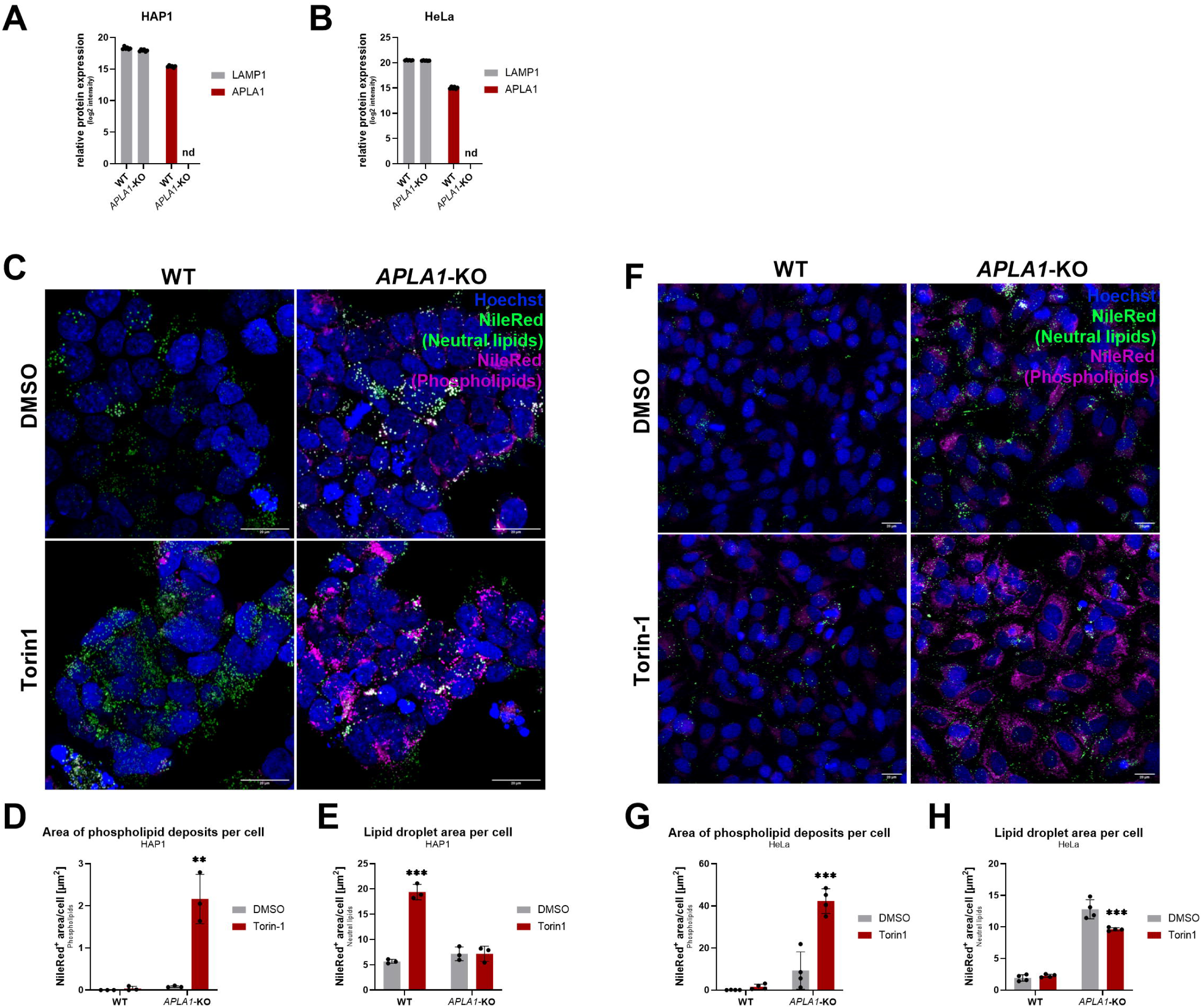
Validation of APLA1 knockout and lipid phenotypes in HAP1 and HeLa cells. (**A, B)** Proteomic confirmation of *APLA1*-deficiency in HAP1 (A) and HeLa (B) cells (n = 6 biological replicates). Peptides corresponding to APLA1 were not detectable (nd), while LAMP1 expression was unchanged. (**C**) Representative confocal microscopy images of HAP1 WT and *APLA1*-KO cells treated with DMSO or 250 nM Torin-1 for 16 h. Cells were stained with NileRed for phospholipids (magenta) and neutral lipids (green) and Hoechst (blue) to detect nuclei (scale bars: 20 µm). (**D, E**) Quantitative analysis of phospholipid deposits (D) and lipid droplets (E) from images acquired in (C). (**F**) Confocal microscopy of HeLa WT and *APLA1*-KO cells and quantification of (**G**) phospholipids and (**H**) lipid droplets. Experiments were performed as described for HAP1 cells (scale bars: 20 µm). Data are represented as mean ± SD (n = 3 fields for HAP1 cells, n = 4 fields for HeLa cells; data representative of two independent experiments for each cell line). Statistically significant differences were determined by two-way ANOVA and corrected for multiple comparisons by Sidák’s post-hoc test (\*\**p*<0.01, \*\*\**p*<0.001).

**Figure S8:**
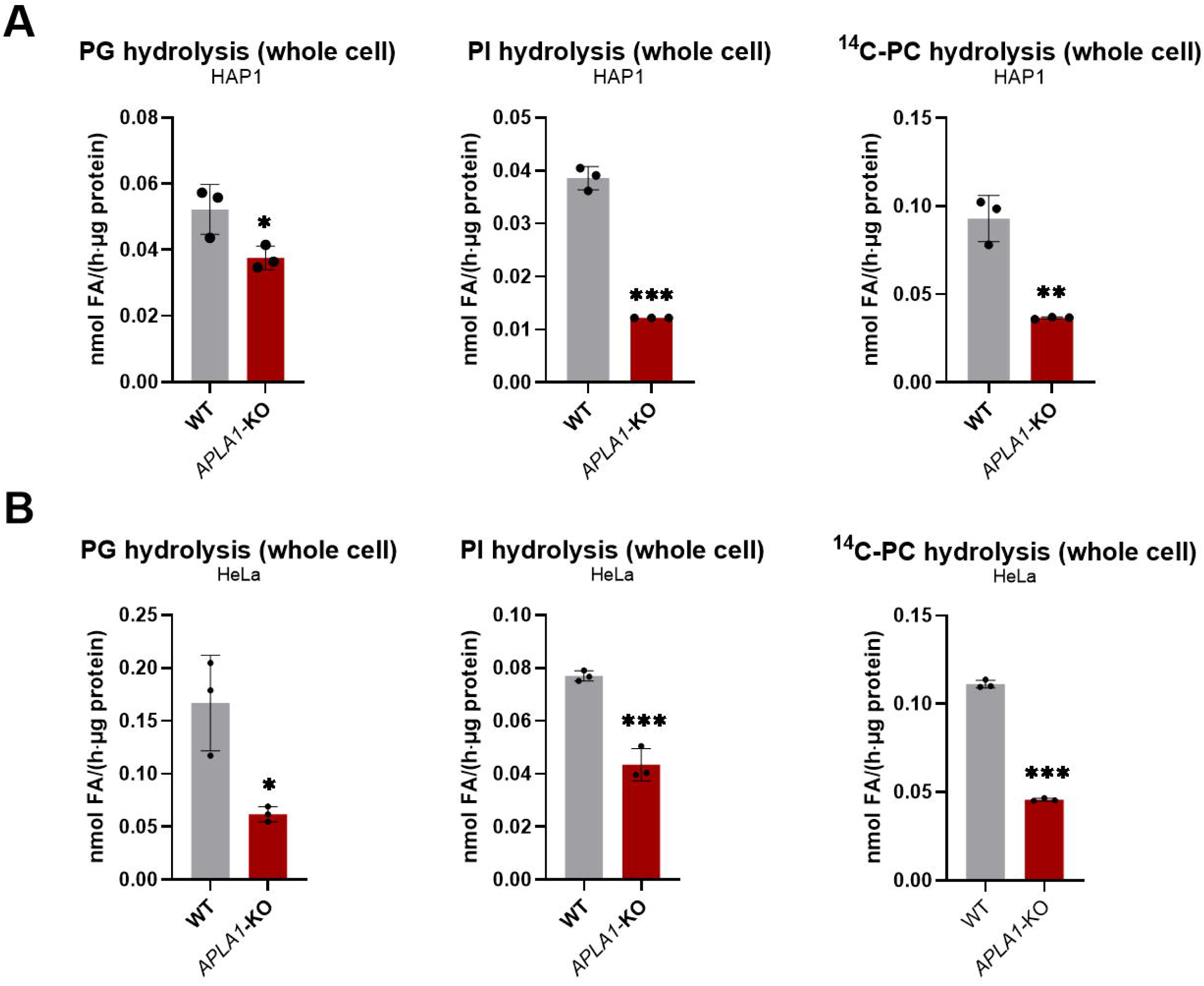
Acid GPL hydrolase activity in whole cell lysates of HAP1 and HeLa WT and *APLA1*-KO cells. Lysates of whole cell fractions (0.67 mg/ml) of (**A**) HAP1 and (**B**) HeLa WT and *APLA1*-KO cells were incubated with 0.33 mM of either PG, PI, or a mixture of PC:BMP (70:30) at pH 4.5 for 1h. For PG and PI hydrolysis, the release of fatty acids was determined using an enzymatic kit. To assess PC hydrolysis, unlabeled PC (233 µM final concentration) was mixed with di-oleoyl-1-^14^C,^14^C-PC (final concentration 6 µM) and unlabeled BMP (30 mol%). The release of ^14^C-oleic acid was quantified by liquid scintillation counting (n = 3 independent replicates). Data are represented as mean ± SD. Statistically significant differences were determined by Students two-tailed unpaired *t*-test (\**p*<0.05, \*\**p*<0.01, \*\*\**p*<0.001).

## Materials and Methods

### Materials

Lipids and internal standards were purchased from Avanti Polar Lipids unless stated otherwise. For storage, the lipid stock solutions were dissolved in CHCl_3_, covered with a layer of N_2_, and kept at # 20°C. All other materials and consumables were purchased form Sigma Aldrich, unless indicated otherwise. HEK293, HEK293T, and HeLa cells were acquired from the American Type Culture Collection (ATCC). HAP1 cells were purchased from Horizon Discovery. Expi293F cells were obtained from Thermo Fisher.

### Reagents, plasmids, and antibodies

All reagents, plasmids, and antibodies used in this study are listed in supplemental tables S1-3.

## Methods

### Cell culture

HEK293 and HeLa cells were grown in DMEM (4.5 g/l glucose; Gibco, Cat#11965084) supplemented with 10% heat-inactivated fetal bovine serum (iFBS), 100 U/mL penicillin, and 100 µg/mL streptomycin (P/S) (= complete media). HAP1 cells were grown in IMDM (Gibco, Cat#12440061) containing 10% iFBS and P/S. For experiments with Torin-1, cells were incubated with media containing either 250 nM Torin-1 (BIOZOL, Cat#SEL-S2827-50MG) or DMSO (vehicle) for 16 h. In some experiments, cells were simultaneously treated with 20 µM amiodarone (Sigma, Cat#A8423-1g). Expi293F cells were grown in Expi293^TM^ expression medium (Gibco, Cat#A1435101). All cells were maintained at 37°C in an environment of 7% CO_2_, and 95% humidified atmosphere. Expi293F cells were cultured in suspension at 125 rpm in Erlenmeyer flasks.

### Generation of KO cell lines

*APLA1*-KO, *PLA2G15*-KO, and dKO cells were generated using CRISPR/Cas9 gene editing. sgRNAs encoding sequences were cloned into the pSpCas9(BB)-2A-Puro (PX459) vector (Addgene, plasmid #62988, *APLA1* sgRNA1: TCTCAGGCGGGAGCATGCTG, *APLA1* sgRNA2: CAGAAGCAGGACCCACGCCG, *PLA2G15* sgRNA: GCTGCTGCTAATGCTGCTCG). Cells were transfected with the constructs using Lipofectamine 3000 (Invitrogen, Cat#L3000001) according to the manufacturer’s instructions. Cells transfected with the empty PX459 vector served as control. Twenty-four hours post transfection, media was changed to media containing puromycin (2 µg/ml) for selection of transfected cells. Cells were then re-seeded into 100 mm dishes at low density and grown until colonies were observed. Single-cell-derived colonies were then picked manually under a microscope. Gene editing was confirmed by Sanger sequencing and loss of proteins was confirmed by proteomic analysis. Control cells, transfected with the empty PX459 vector were subjected to the same procedure. HEK293 dKO cells were generated on the background of *APLA1*-deficient cells.

### Proteomic analysis

Total cell lysates (5% SDS, 50mM AMBIC) were processed using S-Trap™ micro columns (C02-micro-80, Protifi) following the high recovery protocol recommended by the manufacturer and using trypsin (Pierce, Cat#90059) as the carrier protein. Mass spectrometry samples were analyzed on timsTOF ion mobility mass spectrometer (Bruker) in-line a reversed-phase C 18 Aurora column (25 cm × 75 μm) on an UltiMate 3000 UHPLC system (Thermo). The spectra were recorded in DIA mode and processed with DIA-NN (v.1.9) using a synthetic library created from the UniProt protein database (human reviewed, 2024_07_29) with default settings. MS2 and MS1 mass accuracies were set to 20 ppm and the scan window size was set to 9. Output was filtered at 0.01 false discovery rate (FDR).

### Generation of cells stably expressing TMEM192-3xHA for lysosome immunopurification

For stable expression of TMEM192-3xHA, lentiviral particles were generated as follows: HEK293T cells were seeded in 6-well plates. The following day, cells were washed once with PBS before they were incubated for 5 hours with DMEM containing 10% iFBS without P/S, supplemented with 25 µM chloroquine. Cells were then transfected with the psPAX2, pMD2.G, and pLJC5-TMEM192-3xHA plasmids using polyethyleneimine. Cells were incubated overnight before the media was replaced with DMEM containing P/S and 10% iFBS. The virus-containing supernatant was harvested for two consecutive days, combined, cleared by centrifugation, and filtered through a 45 µm filter unit. For infection of HEK293 WT and KO cell lines, cells were seeded in 6-well plates at a density of 1*10^6^ cells/well. The following day, media was replaced with media containing 8 µg/ml polybrene and 0.5 ml of viral suspension before the cells were centrifuged at 300 x g and 32°C for 1 h. 24 h post-infection, infected cells were selected using media containing puromycin. Equal expression of TMEM192-3xHA was confirmed by immunoblotting.

### Stable re-expression of APLA1 and APLA1-S11A in *APLA1*-KO cells

For rescue experiments, HIS-tagged active APLA1 or the inactive APLA1-S111A variant were re-expressed in *APLA1*-deficient cells using a lentivirus. For this, the coding sequence of APLA1 and APLA1-S111A were cloned into the pLVX-IRES-Puro vector. To generate viruses, HEK293T cells were transfected with pLVX-APLA1-HIS-IRES-Puro, psPAX2, and pMD2.G using polyethyleneimine. A virus generated with the empty pLVX-IRES-Puro vector served as an additional empty vector (e.v.) control. Harvesting of viruses and infection of cells were performed as described above for stable expression of TMEM192-3xHA.

### Microscopy studies

#### LysoTracker staining

Cells were seeded into poly-d-lysine-coated 35 mm glass bottom dishes (Ibidi, Cat#81218-200) at a density of 500,000 cells/well. The following day, the media was aspirated and replaced with media containing 1 µg/ml Hoechst 33342 (Abcam, Cat#228551) and 60 nM LysoTracker Red DND-99 (ThermoFisher, Cat#L7528). After incubation for 1 h, the media was replaced with standard growth media and cells were immediately imaged.

#### NileRed staining, LysoSensor costaining and fluorescent fatty acid tracing

For NileRed staining, cells were seeded in poly-d-lysine coated 8-well chamber slides (SARSTED, Cat# 94.6190.802) at a density of 100,000 cells/well. The next day, cells were treated with either 250 nM Torin-1 or DMSO for 16 h before they were fixed for 20 minutes using a 4% PBS-buffered formaldehyde solution containing 1 µg/ml Hoechst 33342 (Abcam, Cat#228551). Fixed cells were washed with PBS before they were incubated with PBS containing 1 µg/ml NileRed (ThermoFisher, Cat#415711000). After 15 minutes, the staining solution was replaced with fresh PBS and cells were imaged using a confocal microscope. Colocalization studies with LysoSensor were performed in live cells. For this, living cells were stained with 1 µM LysoSensor Green DND-189 (Invitrogen, Cat#L7535) for 1 h before they were stained with Hoechst (1 µg/ml) and NileRed (1 µg/ml) for an additional 15 minutes and image acquisition was performed.

For tracing of fluorescently labeled fatty acids, cells were seeded as described above before they were pulsed with 4 µM BODIPY-labeled fatty acids (BODIPY FL C16, Invitrogen, Cat# D3821). After 5 h of incubation, cells were washed twice with media before the media was changed to complete media containing either 250 nM Torin-1 or DMSO for 16 h. Cells were fixed and stained with NileRed as described above and analyzed by confocal microscopy.

#### Tracing of fluorescent PG

To assess PG accumulations, HEK293 WT and KO cells were seeded in poly-d-lysine coated 8-well chamber slides as described above. The next day, cells were treated with 2 µM 18:1-12:0 NBD PG (Avanti, Cat#A81166) for 4 h. Subsequently, cells were washed twice and media containing either 250 nM Torin-1 or DMSO (vehicle) was added. After 16 h incubation, cells were washed and fixed for confocal microscopy using 4% PBS-buffered formaldehyde.

### Image acquisition and analysis

Imaging was performed on a Leica SP8 confocal laser scanning microscope equipped with a HC PL APO CS 63×/1.2 NA water immersion objective. Care was taken to ensure uniform treatment of cells across all conditions and consistent imaging settings were applied throughout all samples in individual experiments.

For image analysis the open-source software Fiji was used. For quantification of LysoTracker and NileRed signals, channels were split and a maximum intensity projection was generated for each channel. Background subtraction was performed using a rolling ball radius of 50 pixels. Threshold values were individually set for each channel and kept constant for all images throughout all conditions in each experiment. Based on the threshold, a mask was created and the “Watershed” algorithm was applied before particles were analyzed using the in-built “Analyze particle” function, with a particle size ranging from 0 to infinity for LysoTracker^+^/NileRed^+^ structures. NileRed^+^ lipid droplets were subtracted from the NileRed phospholipid channel to account for residual crosstalk and bleed-through. To determine the average LysoTracker^+^/NileRed^+^ area/per cell, positive areas were summed and divided by the number of nuclei on the respective image. The number of cells in each projection was determined by manual counting of nuclei detected using Hoechst nuclear stain. For colocalization studies of NileRed with LysoSensor and BODIPY FL C16, signal intensities of both channels across the whole image (LysoSensor) or along a linear region (BODIPY FL C16) were plotted against the ROI distance/area.

For improved visualization, brightness and contrast values were adjusted (same settings were applied to all images throughout all conditions).

### Electron microscopy

Cells were seeded in 6-well plates on Aclar film (Gröpl), at a density of 1*10^6^ cells/per well before they were treated with 250 nM Torin-1 for 16 h. Cells were fixed in 2.5% (w/v) glutaraldehyde and 2% paraformaldehyde (w/v) in 0.1 M cacodylate buffer, pH 7.4, for 3 h, and then post-fixed in 2% (w/v) osmium tetroxide for 2 h at room temperature. After dehydration (in graded series of ethanol), cells were infiltrated in propylene oxide and TAAB Embedding Resin overnight, afterwards in pure TAAB Embedding Resin (3 h), (TAAB Laboratories Equipment Ltd., UK) transferred into embedding moulds, and polymerized (48 h, 60°C). Ultrathin sections (70 nm) were cut with a UC 7 Ultramicrotome (Leica Microsystems, Austria) and stained with platinum blue (EMS, USA) for 15 min and lead citrate for 5 min.

For scanning transmission electron microscopy (STEM), imaging mode of a field emission scanning electron microscope (ZEISS FE-SEM Sigma 500) with an acceleration voltage of 15 kV in combination with ATLAS TM was used to correlate conventional TEM and STEM images, on large areas with high-resolution AZoNano^36^. The diameter of lysosomes/multilamellar organelles were determined using Fiji.

### Tracing of radiolabeled fatty acids

For tracing of ^3^H-labeled fatty acids, cells were seeded in poly-d-lysine coated 12-well plates at a density of 500,000 cells per well. The next day, cells were pulsed with 25 nM ^3^H-oleic acid (9,10-^3^H oleic acid). After 5 h, cells were washed twice with complete media before complete media containing either 250 nM Torin-1 or DMSO (vehicle) was added. After 16 h, cells were washed with PBS before lipids were extracted directly from the adherent cells by adding 500 µl hexane:2-propanol (3:2 v/v). The plate was incubated on a shaker for 5 minutes before the extraction mixes were collected in 2 ml safe lock tubes. The extraction was carried out a second time, followed by a final extraction step with 400 µl of 2-propanol. Respective extracts were combined and dried under a stream of N_2_. The dried lipid extracts were reconstituted in 500 µl of chloroform. An aliquot of each lipid extract was counted by liquid scintillation to assess tracer uptake into the total lipid pools. The remaining extracts were spiked with 10 µl of a 1 mg/ml TAG standard solution to allow visualization of thin-layer chromatography (TLC)-separated lipid bands in iodine-vapor. Solvents were evaporated under a stream of N_2_ before the dried lipid extracts were reconstituted in chloroform and spotted onto a silica TLC plate. Lipids were separated using hexane:diethylether:acetic acid (70:29:1 v:v:v) as mobile phase. TAG bands were cut from the silica plate and the comigrating radioactivity was determined by liquid scintillation counting.

### Lysosome immunopurification (LysoIP)

Lysosomes were purified by LysoIP as previously described by others^18,37^. In brief, HEK293 WT and KO cells stably expressing TMEM192-3xHA were seeded in 145 mm dishes and grown until ∼90% confluency. Cells were then washed twice with ice-cold PBS before they were harvested in ice-cold KPBS (136 mM KCl, 10 mM KH_2_PO_4_, pH 7.25) and centrifuged at 1,000 x g and 4°C for 2 min. The supernatant was removed, the pellet resuspended in KPBS, and an aliquot of the cell suspension was taken for whole cell analysis. The remaining cells were gently lysed by repeated strokes through a 26G needle using 1 ml syringes. The lysate was centrifuged at 1,000 x g and 4°C for 2 min and the supernatant was transferred to tubes containing 80 µl of pre-washed Pierce^TM^ anti-HA magnetic beads (Thermo Scientific, Cat#88837). Lysosomes were allowed to bind for 4 min before tubes were placed on a magnetic rack and beads were allowed to be pulled by the magnet. The residual lysate was removed before the beads were washed in 3 consecutive steps with KPBS. After the last wash, the supernatant was removed and lysosomes bound to beads were resuspended in 1 ml of KBPS and transferred to fresh tubes, the supernatant was removed and lysosomes bound to beads were snap frozen in liquid N_2_ and stored for further processing at #80°C.

To generate lysosomal lysates for activity assays, lysosomes were lysed by inducing a hypotonic shock after the last wash. For this, lysosomes bound to beads were resuspended in 100 µl of ddH_2_O and incubated on ice for 1 h before beads were removed using a magnetic rack. The lysate was then centrifuged at 16,000 x g for 10 minutes to remove residual debris. The supernatant was transferred to fresh tubes and used for enzymatic assays. Protein concentration of lysates was determined by Bradford protein assay (Bio-Rad, Cat#5000001).

### Transient transfection of cells

For enzymatic activity assays, cells were transfected with plasmids encoding β-galactosidase (LacZ), human APLA1, murine APLA1, human PLA2G15, or human APLA1-S111A using Lipofectamine 3000 according to the manufacturer’s instructions. In brief, cells were seeded in 6-well plates or 100 mm dishes and grown until 80-90% confluency. Cells were then transfected with protein-encoding plasmids and maintained for an additional 24-48 h before harvesting.

### Preparation of cell lysates

Cells were washed twice with PBS before they were harvested by scraping in ice-cold PBS. Cell suspensions were centrifuged at 300 x g and 4°C for 3 min before the supernatant was removed and the pellet resuspended in lysis solution (0.25 M sucrose, 1 mM EDTA, 1 mM dithiothreitol, 20 µg/ml leupeptin, 2 µg/ml antipain, 1 µg/ml pepstatin, pH 7). Cells were disrupted by sonication on ice using a SONOPULS ultrasonic homogenizer (Bandelin, Germany) equipped with a T72 sonotrode for 3×10 sec with an amplitude of 15%. The lysate was centrifuged at 1,000 x g and 4°C for 10 min to remove cellular debris and nuclei. Protein concentrations of lysates were determined by Bradford protein assay (Bio-Rad, Cat#5000001).

### Lipid substrate preparation

Lipid substrate solutions were prepared in PBS with a final lipid concentration of 1 mM unless stated otherwise. For this, the required volume of lipid dissolved in CHCl_3_ was dried under a stream of N_2_. The dried lipids were then reconstituted in PBS by sonication using a SONOPLUS ultrasonic homogenizer (Bandelin) with 15% amplitude for 10 seconds.

### In vitro assays with lysates

For in vitro assays with whole cell lysates, 20 µl of lysate (2 mg protein/ml) were mixed with 20 µl of the indicated buffer (0.1 M) and 20 µl of lipid substrate solution (1 mM), unless indicated otherwise. The reactions were then incubated at 400 rpm and 37°C for 1 h on a thermomixer. For experiments with isolated lysosomes, lysates of purified organelles (0.4-1 µg protein) were incubated with substrates and buffer as described above. To determine PLB and PLC activity in purified lysosomes, lysates of WT and dKO cells were incubated with substrate and buffer in the presence or absence of 1 µM of the lysosomal acid lipase (LAL) inhibitor Lalistat2 (LALi2) (Sigma, Cat#SML2053) and reaction products (diacylglycerols, ceramides, and lyso-GPLs) were determined by targeted MS analysis. For measurement of non-esterified fatty acids (NEFA), reactions were performed in 96-well plates. Assays and subsequent lipid extraction for further analysis by thin layer chromatography (TLC) or mass spectrometry were performed in 2 ml safe-lock Eppendorf tubes. For experiments with CADs, the substrate/buffer mix was premixed with the respective compound before adding it to the reaction mix.

### Measurement of non-esterified fatty acids (NEFA)

Free fatty acids generated by enzymatic activities from various phospholipid substrates were determined using the NEFA-HR(2) assay kit (FUJIFILM Wako, NEFA-HR(2) R1: 434-91795, NEFA-HR(2) R2: 436-91995, NEFA Standard: 270-77000) according to the manufacturer’s instructions. In brief, reactions performed in 96-well plates were incubated with 100 µl NEFA-HR(2) R1 for 10 min at 37°C before 50 µl of NEFA-HR(2) R2 were added. The reactions were incubated for another 5 min at 37°C before measuring the absorbance at 560 nm. A serial dilution of the NEFA Standard was used to quantitatively determine liberated fatty acids.

### Screening for acid phospholipases

To screen for acid phospholipase activities, a previously established library of murine lipid hydrolases and related enzymes was utilized^11^. Therefore, 4 µl of lysates (2 mg/ml) of enzyme-expressing Expi293F cells were incubated with 4 µl substrate solution (2 mM lipid emulsified either in 0.1 M citrate buffer (pH 4.5) or 0.1 M Tris-HCl buffer (pH 7.4)) containing either PG, PI, or PS at 37°C in a 384-well plate for 1 h. The release of fatty acids was determined using the NEFA assay kit as described above (20 µl NEFA-HR(2) R1 and 10 µl NEFA-HR(2) R2). Activities exceeding a z-value of +1 were considered positive hits. Localization of potential candidates was determined using the UniProt database^38^.

### Site-directed mutagenesis

To generate a catalytically inactive mutant of APLA1, the coding sequence of the active site serine (S111) was mutated to an alanine (S111A). Mutagenesis was performed using the Q5 site-directed mutagenesis kit (NEB, Cat#E0552S) according to the manufacturer’s instructions (primers: APLA1-S111A-fw: CATCTGCTACGCGCAGGGGGGCC, APLA1-S111A-rev: AGATGCACCCCTTGAGGGGC. Annealing temperatures were set according to the NEB primer tool. Elongation time was set to 3:30 min (∼20-30 sec/kb). Mutagenesis was verified by Sanger sequencing.

### Recombinant expression and purification of APLA1 and APLA1-S111A

HIS-tagged APLA1 and APLA1-S111A were expressed using the Expi293 system according to the manufacturer’s instructions. In brief, cells were grown in 25 ml media in 125 ml vented Erlenmeyer flasks (Corning, Cat#431143) to a density of 3*10^6^ cells/ml and subsequently transfected with plasmids encoding HIS-tagged proteins using the ExpiFectamine^TM^293 reagent (Gibco, Invitrogen GmbH, Cat#A14524) according to the manufacturer’s instructions. 24h post-transfection, Transfection Enhancer 1 and ExpiFectamine^TM^ 293 Transfection Enhancer 2 were added to the cells. 48h post-transfection, cells were pelleted by centrifugation and the supernatant (SN) was collected. The SN was diluted 1:1 with PBS and incubated with 250 µl of pre-washed PureCube 100 Ni-INDIGO agarose (Cube Biotech, Cat#75103). HIS-tagged proteins were allowed to bind overnight at 4°C on an over-top shaker. The following day, the resin was centrifuged at 300 x g for 2 min and the SN was removed. The resin was resuspended in 500 µl of buffer W1 (50 mM HEPES pH 7.25, 500 mM NaCl, 0.1 mM EDTA, 5 mM beta-mercaptoethanol, 5% glycerol, 20 mM imidazole, 1 mM DTT, 1% Triton X-100, 20 µg/ml leupeptin, 2 µg/ml antipain, 1 µg/ml pepstatin, pH 7.25) and loaded onto a gravity column (BioRad, Cat#7311550). The beads were then washed in four consecutive wash steps with the following buffers: wash 1 (buffer W1), wash 2 (buffer W2, 50 mM HEPES pH 7.25, 500 mM NaCl, 0.1 mM EDTA, 5 mM beta-mercaptoethanol, 5% glycerol, 10 mM imidazole, 1 mM DTT, 20 µg/ml leupeptin, 2 µg/ml antipain, 1 µg/ml pepstatin, pH 7.25), wash 3 (buffer W3, 50 mM HEPES pH 7.25, 250 mM NaCl, 0.1 mM EDTA, 5 mM beta-mercaptoethanol, 5% glycerol, 1 mM DTT, 20 µg/ml leupeptin, 2 µg/ml antipain, 1 µg/ml pepstatin, pH 7.25), and wash 4 (buffer W4, 50 mM HEPES pH 7.25, 125 mM NaCl, 0.1 mM EDTA, 5 mM beta-mercaptoethanol, 5% glycerol, 1 mM DTT, 20 µg/ml leupeptin, 2 µg/ml antipain, 1 µg/ml pepstatin, pH 7.25). Proteins were eluted in fractions using elution buffer (50 mM HEPES pH 7.25, 125 mM NaCl, 0.1 mM EDTA, 5 mM beta-mercaptoethanol, 5% glycerol, 250 mM imidazole, 1 mM DTT, 20 µg/ml leupeptin, 2 µg/ml antipain, 1 µg/ml pepstatin, pH 7.25). Protein-containing fractions were pooled, dialyzed twice against PBS containing 1 mM EDTA and 20 µM DTT, and snap frozen in liquid N_2_.

### SDS-PAGE and immunoblotting

For the analysis of proteins and cell lysates samples were mixed with SDS-sample buffer and heated to 95°C for 10 min prior to loading them on a 10% SDS-polyacrylamide gel. Proteins were separated by applying a constant current of 25 mA. Gels were then either stained using Coomassie brilliant blue solution or used for immunoblotting. For this, proteins were transferred onto a polyvinylidene fluoride (PVDF) membrane using a blotting chamber filled with CAPS buffer (10 mM CAPS, 10% MeOH (v/v), pH 11) and a constant current of 200 mA for 70 min. The membranes were blocked for 1 h with 10% blotting grade milk powder dissolved in TST (50 mM Tris/HCl, pH 7.4, 0.15 M NaCl, 0.1% Tween-20). Membranes were incubated for either 2 h (RT) or overnight (4°C) with primary antibodies dissolved in TST containing 5% milk powder. Secondary antibodies were prepared in TST containing 5% milk powder. Bands were visualized and detected using Clarity Western ECL Substrate (Bio-Rad, Cat#1705060) and the ChemiDoc Touch Imaging System (Bio-Rad). All antibodies used in this study are listed in supplemental Table S3.

### Surface charge prediction

The PDB-file of human PPT2 (PDB-ID: pdb_00001pja) was prepared with PDB2PQR at pH values of 5.5, 6.0, and 6.5 with titration states and charges assigned and hydrogens added. The electrostatic properties were generated using ABPS. Both steps were performed using default parameters in the PDB2PQR and ABPS biomolecular solvation software suite^39^. Images were generated using PyMOL Molecular Graphics System (version 2.4.1, Schrödinger LLC).

### Molecular dynamics and computational analysis

#### Template crystal structure

The starting coordinates for PPT2 (APLA1) were taken from the crystal structure deposited in the Protein Data Bank under accession code pdb_00001pja. This structure resolves the mature APLA1 hydrolase fold, and defines the catalytic Ser–His–Asp triad that is responsible for the enzyme’s lipid hydrolase activity. The deposited model was used as the input template for subsequent system building. Crystallographic waters and non-biological additives were removed, and the biological protomer was retained as the single protein chain (PROA) for membrane insertion. Because APLA1 acts at the lumenal face of the lysosomal membrane, the protein was treated as a peripheral, membrane-associated species rather than an integral membrane protein.

#### Preparation of protein-membrane systems

Protein–membrane complexes were assembled with the Membrane Builder module of CHARMM-GUI^40,41^ using the CHARMM36m all-atom protein force field^42^ together with the CHARMM36 lipid force field^43^ and the TIP3P water model^44^ throughout. Prior to bilayer assembly, APLA1 was oriented relative to the membrane normal with the PPM/OPM procedure^45^ so that the surface loops flanking the active-site entrance faced the lipid headgroup region; this orientation was then held fixed while the surrounding bilayer was packed around the protein. A rectangular bilayer patch was generated on either side of the protein, hydrated with an approximately 60 Å water slab above and below the membrane, and neutralized in 0.15 M NaCl (Na^+^/Cl^−^) using the Monte-Carlo ion-placement scheme. All titratable residues (acidic, basic, and His) were built in their standard CHARMM protonation states at this stage. To reproduce the acidic milieu of the lysosomal lumen, the protein was propagated under constant-pH molecular dynamics (cpHMD) at a simulation pH of 4.5, using the scalable λ-dynamics constant-pH implementation available in the constant-pH branch of GROMACS7 together with the CHARMM36m constant-pH force-field port (charmm36-mar2019-cphmd). CHARMM-GUI outputs were carried forward into production and defined the two systems:

PC (single-component bilayer). A pure phosphatidylcholine (PC) bilayer comprising 768 PC molecules (384 per leaflet), with the APLA1 protomer. The system was solvated with 132,200 TIP3P water molecules and neutralized with 375 Na^+^ and 365 Cl^−^ ions in a periodic box of approximately 164.9 × 164.9 × 196.3 Å^3^.

PC/BMP (anionic mixed bilayer). A two-component bilayer at a 1:1 molar ratio of PC to bis(monoacylglycero)phosphate (BMP), built with 384 PC and 384 BMP molecules (modeled with the CHARMM BMGP residue) for a total of 768 lipids. BMP carries a net −1 charge per molecule, so the mixed bilayer was neutralized with a correspondingly larger sodium count (745 Na+ and 351 Cl−) in addition to the 0.15 M background salt, in a periodic box of approximately 162.2 × 162.2 × 196.3 Å3 containing 127,124 TIP3P waters. This composition was chosen to emulate the anionic, BMP-enriched intralysosomal membranes on which APLA1 is active.

The system was prepared for constant-pH simulations using the phbuilder tool (https://gitlab.com/gromacs-constantph/phbuilder). All acidic and basic side chains of APLA1 were made titratable: 60 titratable sites in total, spanning aspartate (ASPT, reference pKa 3.65), glutamate (GLUT, 4.25), histidine (HSPT, represented as a coupled three-state model comprising the two neutral δ/ε tautomers and the doubly protonated form), lysine (LYST, 10.4) and arginine (ARGT, 13.8). Because protonation and deprotonation change the total charge of the system, electroneutrality was preserved on-the-fly by coupling all sites to a collective titrating buffer (BUF) through a charge constraint, so that any charge released by a titrating residue was compensated by an equal and opposite change distributed over the buffer particles (buffer multiplier 121); histidine’s three states were additionally handled with a multistate constraint. This yielded 61 λ-atom collections in total. Following the recommended settings for accuracy and sampling, each λ particle was assigned a mass of 5.0 (in the reduced GROMACS λ units) and evolved with a coupling time constant of 2.0 ps under a double-well barrier of 7.5 kJ mol−1, with the λ coordinates updated every 500 steps. Holding the simulation at pH 4.5 allowed the ionizable residues to dynamically sample their lysosome-relevant protonation states throughout the trajectories, and constant-pH dynamics were maintained throughout equilibration, production, steered MD, and umbrella sampling.

#### Molecular dynamics simulations

All simulations were performed using the constant-pH build of GROMACS. Non-bonded interactions followed the CHARMM-recommended settings: a Verlet neighbor-search scheme, a 1.2 nm real-space cut off for both electrostatics and van der Waals interactions, a force-switch modifier applied to the Lennard-Jones potential between 1.0 and 1.2 nm, and long-range electrostatics evaluated by particle-mesh Ewald (PME)^46^ with a 0.14 nm Fourier grid spacing. Bonds to hydrogen were constrained with the LINCS algorithm^47^ (order 4), permitting a 2 fs integration time step.

Each system was first energy-minimized by steepest descent (up to 5000 steps, until the maximum force fell below 1000 kJ mol^−1^ nm^−1^) and then equilibrated in stages with harmonic restraints applied to the protein, the lipid head-group heavy atoms (force constant 1000 kJ mol^−1^ nm^−2^) and to the lipid dihedral angles, following the CHARMM-GUI protocol. Equilibration comprised a restrained NVT stage followed by a restrained NPT stage, after which the restraints were released for production. Production dynamics were propagated in the NPT ensemble in successive 100 ns segments (50,000,000 steps at 2 fs). Temperature was maintained at 303.15 K with the velocity-rescale thermostat12 (τ = 1.0 ps) using three separate coupling groups for the solute, membrane, and solvent, and the pressure was held at 1.0 bar with the C-rescale barostat^48^ applied semi-isotropically (τ = 5.0 ps, compressibility 4.5 × 10−5 bar^−1^) to allow the membrane area to relax independently of the box height. Each system was run for 500 ns of production (five consecutive 100 ns segments). For both membrane compositions, the last frame of this 500 ns production run was centered in the box and used as the starting configuration for the steered-MD stage.

#### Umbrella sampling and free-energy analysis

Umbrella-sampling windows were selected from the extracted SMD frames so as to give an approximately uniform spacing of ∼0.1 nm along the protein–membrane COM reaction coordinate, from the membrane-bound contact state out to full separation in bulk water, spanning COM distances of roughly 1.78–5.0 nm. In each window, the protein was restrained at its target separation by a harmonic umbrella potential (force constant 400 kJ mol^−1^ nm^−2^) acting on the z-component of the protein–phosphate COM distance, using the same cylindrical reference geometry (25 Å radius) as in the SMD stage and referencing the upper-leaflet phosphate group. Each window was first equilibrated for 1 ns in the NPT ensemble (500,000 steps) and then sampled for 20 ns of production umbrella dynamics (10,000,000 steps at 2 fs), again under constant-pH conditions at pH 4.5. The biased distributions from all windows were unbiased and combined using the weighted histogram analysis method (WHAM) as implemented in gmx wham^49^ over the range 1.78–5.0 nm. Statistical uncertainties on the potential of mean force were estimated by Bayesian bootstrapping of complete histograms (200 bootstrap resamples). The resulting free-energy profiles were referenced to the membrane-bound minimum, so that the depth of the well relative to the dissociated plateau reports the free energy of association of APLA1 with each bilayer. Profiles were computed independently for the PC and PC+BMP systems and are reported in kcal*mol^−1^. ChimeraX (version 1.11.1) was used for visualization.

#### Covalent APLA1–phospholipid complex prediction

Near-attack protein–ligand conformations were predicted directly with the deep-learning co-folding model Boltz-2^50^. The mature APLA1 sequence (UniProt: Q9UMR5, with its N-terminal signal peptide removed) was folded together with each phospholipid ligand under an explicit covalent-bond restraint linking the Oγ atom of the catalytic serine (S111; the nucleophile of the G-X-S-X-G elbow) to the carbonyl carbon of the reactive acyl chain, bringing the two reactive atoms into contact so as to capture the near-attack conformation. The crystal structure was supplied as a folding template, and a precomputed multiple-sequence alignment was reused across all jobs; five models were sampled per complex.

Because diacyl phospholipids present two ester carbonyls to the nucleophile, each lipid was modeled in both covalent regiochemistry: attachment through the *sn-1* acyl carbonyl and, separately, through the *sn-2* acyl carbonyl.

All of the near-attack APLA1–phospholipid complexes were predicted with high confidence (range 0.92–0.95 across all headgroups and both regiochemistry). The extent of each near-attack interface was quantified by the covalent-ligand interface-atom pairs count (cLIGPA) score, which was adopted and modified from local interaction area (cLIA)^51^, the number of protein–ligand atom pairs in direct contact at the modeled interface (Cβ–ligand-atom distance ≤ 8 Å with a predicted aligned error ≤ 12 Å). cLIGPA was taken from the single most confidently docked pose: for each complex the model that maximizes the interface-confidence score iLIS^51^ was selected, and its cLIGPA reported. PyMOL (version 3.0) was used for visualization.

### In vitro assays with purified APLA1 and APLA1-S111A

For in vitro assays, purified proteins at the indicated concentrations (0.2 – 2 µg) were incubated with 20 µl of 1 mM lipid substrate solution and 30 µl of buffer solutions at 37°C and 400 rpm on a thermomixer for 30-60 min. Assays were performed in 96-well plates. Activity was determined by measuring non-esterified fatty acids (NEFA) as described above for assays with cell lysates. To investigate the positional preference of APLA1, assays were performed in 2 ml safe lock Eppendorf tubes and lipids were extracted. Lyso-GPLs (LPG 18:1, LPG 18:0, LPC 16:0, or LPC 20:4) generated from phospholipid substrates in experiments to determine the positional preference of APLA1 were detected by targeted MS-analysis. For experiments with cationic amphiphilic drugs (CADs, amiodarone or chloroquine, Sigma Aldrich, Cat#C6628), substrate and buffer solutions were premixed and pre-incubated for 5 min with CADs at the indicated concentrations before they were added to the purified enzyme.

### Liposome preparation for liposome pulldown experiments

PC and PC/BMP liposomes were prepared as follows: Lipids in indicated ratios were dried under a stream of N_2_. The dried lipids were then hydrated in PBS for 30 min at RT before the suspension was briefly vortexed. Unilamellar liposomes with a diameter of 100 nm were then generated by repeated extrusion (15x) through a 0.1 µm polycarbonate membrane using a mini extruder (Avanti, Cat#610000-1EA).

### Liposome pulldown

For pulldown experiments, 400 µl of 1 mM PC or PC/BMP (70/30 mol%) liposome solutions were filled up to a final volume of 1 ml with PBS and then pelleted by centrifugation at 100,000 x g and 22°C for 30 min. The SN was removed and the pellets resuspended in 200 µl of the indicated buffer to reach a final lipid concentration of 2 mM. The liposomes solutions were then incubated with 4 µg of purified APLA1 in a final volume of 75 µl. The solutions were incubated for 10 min at 37°C before the mix was filled up with 1 ml of mineral oil and centrifuged at 100,000 x g and 22°C for 30 min. The oil was removed and the remaining SN was transferred to fresh Eppendorf tubes and mixed with 25 µl of 4x SDS sample buffer. Pellet fractions were directly resuspended in 100 µl of 1x SDS sample buffer. 35 µl of each fraction were subsequently loaded onto an SDS gel and an SDS-PAGE was performed. APLA1 was visualized by staining the gel with Coomassie brilliant blue. For experiments with cationic amphiphilic drugs (CADs), liposomes were pre-incubated with amiodarone (final concentration 300 µM) or DMSO (vehicle) at 37°C for 10 minutes before APLA1 was added.

### Lipid extraction

Lipids from samples were extracted according to Matyash et al. with modifications^52^. Samples were collected in 2 ml safe lock Eppendorf tubes and resuspended in 700 µl of methyl-tert-butylether (MTBE)/MeOH (3:1; v/v). For LC-MS measurements, internal standards were added beforehand to the extraction mix (supplementary table S1). Samples were extracted under constant shaking on a thermomixer at 1,100 rpm for 20 min at RT before 140 µl ddH_2_O were added. Samples were incubated for another 10 minutes under constant shaking before they were centrifuged to achieve phase separation at 14,000 × g and RT for 5 min. The upper organic phase was collected, dried under a stream of N_2_, and reconstituted in MeOH/2-propanol/ddH_2_O (6:3:1; v/v/v) for UPLC-MS analysis or CHCl_3_ for thin layer chromatography.

### Thin layer chromatography

Dried lipid extracts were reconstituted in 2x 20 µl CHCl_3_ and spotted onto a TLC Silica gel 60 20 x 20 cm aluminum sheet (Merck, Cat#105553.0001). Lipids were separated using CHCl_3_/MeOH/H_2_O (65:35:5, v/v/v) as mobile phase. The silica plate was immersed in charring solution (25% EtOH, 8.5% H_3_PO_4_; 5% CuSO_4,_ v/v/w) and lipids were visualized by charring at 130°C. Images of plates were captured using the ChemiDoc Touch Imaging System (Bio-Rad). Densitometric analysis of lipid bands was performed using the Image Lab software (Bio-Rad). Data are expressed as arbitrary units (AU). To enhance visualization, brightness and contrast were adjusted.

### MS analysis

#### Lipidomic analysis by UHPLC-QTOF-MS/MS

Samples were analyzed on a UHPLC 1290 Infinity II system coupled to a 6560 QTOF mass spectrometer (Agilent, Waldbronn, Germany). Chromatographic separation was performed on a BEH C18 column (2.1 × 150 mm, 1.7 μm; Waters, USA) at 50°C and 200 μl/min, using (A) water and (B) 2-propanol, both containing 0.1 % formic acid, 10 mmol/l ammonium acetate, and 7.7 µmol/l phosphoric acid. The gradient started at 60 % A (held 0–0.5 min), increased linearly to 20 % A by 9.0 min, and to 0% A by 22.0 min, held until 24.5 min, then returned to initial conditions at 25.0 min and re-equilibrated until 30.0 min. 1 µl of each sample was injected in positive mode, and 5 µl in negative mode.

Detection was performed in TOF-only mode using a dual AJS ESI source with the following parameters: gas temperature 300 °C, drying gas flow 10 L/min, nebulizer pressure 50 psig, sheath gas temperature 400 °C, sheath gas flow 12 L/min, capillary voltage 3500 V, nozzle voltage 500 V, fragmentor voltage 300 V, skimmer1 voltage 30 V, and octopole RF peak 750 V. Continuous mass axis calibration was performed using reference masses of 322.0481 and 922.0098 m/z in positive mode, and 965.9998 m/z in negative mode, with identical source parameters applied in both modes.

Data were acquired in data-dependent acquisition (DDA) mode. A subset of the data was annotated using MS-DIAL^53^ to build a target list, which was subsequently used to annotate the full dataset in Skyline^54^. Peak integrations were manually reviewed and curated based on the expected retention time order, as predicted by the equivalent carbon number^55^. Data were exported and normalized to class-specific internal standards, and total protein content was determined using the Pierce BCA Protein assay.

#### Targeted MS analysis

For targeted analysis of reaction products from in vitro assays with lysosomal lysates and purified proteins (PLB and PLC activity in lysosomal lysates and the positional preference of purified APLA1) and the analysis of BMP, LPG, and PG by mass spectrometry, total lipids were extracted and reconstituted as described above. Chromatographic separation of lipids was performed using an Agilent 1290 Infinity II UHPLC equipped with an ACQUITY UPLC BEH C18 column 2.1 × 150 mm, 1.7 µm (Waters Corporation, Cat#186002353), at a flow rate of 0.2 ml/min, an injection volume of 2 μl, and a column temperature of 50 °C. A 30 min gradient elution was performed, composed of mobile phase A (MeOH/H2O (8/2, v/v)) and B (2-propanol/MeOH (8/2, v/v)), both containing 10 mM ammonium acetate, 0.1% formic acid, and 8 µM phosphoric acid. The gradient started with a linear increase of 50% to 60% mobile phase B over 13 min, followed by a 7 min linear gradient to 100% mobile phase B which was held for 5 min before returning to 50% mobile phase B for 5 min. Lipid species were detected on an Agilent 6470 triple-quadrupole mass spectrometer with Agilent Jet Stream ESI (Agilent Technologies) controlled by Agilent MassHunter Acquisition software version 10.1. Mass spectrometry was performed in fast polarity-switching mode, enabling the acquisition of positive# and negative-ion data within a single run.

Data processing was performed using the Skyline MS analysis software V26.1 and Agilent MassHunter qualitative analysis software V10.1. Data were normalized for recovery, extraction, and ionization efficacy by calculating analyte/internal standard ratios (AU). External one-point calibration was performed for quantitation of in vitro assays. All transitions with respective settings are listed in supplementary tables S4 and S5.

## Data visualization and statistical analysis

Data visualization and statistical analysis were performed using Prism version 10.5.1 except for Fig 2J for which RStudio (version 2026.06.0) was used. Data are presented as mean ± standard deviation (SD) unless stated otherwise. The number of replicates and the statistical tests applied are reported in the figure legends. Significance levels are shown as *p*-values in the figures. ChemDraw version 23.1.1 was used to visualize chemical structures in Fig. 4A, and 5E, F. BioRender was used to create the figure schematics in Fig. 1A. CorelDraw version 23.1.0.389 was used to create schematics and illustrations in Figure 4A.

## Resource availability

### Lead contact

Additional information, inquiries and requests for all reagents and resources should be directed to and will be fulfilled by the lead contact, Robert Zimmermann.

### Materials availability

All unique reagent (e.g. plasmids) generated in this study are available from the lead contact without restriction.

### Data and code availability

Any additional information required to reanalyze the data in this paper is available from the lead contact upon request.

## Supplemental information

Supplemental information is available for this paper. The supplementary data file contains tables with information on chemicals, plasmids, antibodies, and targeted MS precursor-to-product transitions (supplementary tables).

## Author contributions

M.T., D.K., A.L. and R.Z. conceptualized the study. M.T., J.B., M.O., U.T., and N.F. developed the methodology. M.T., J.B., C.Z., D.B., A.P., C.W., A.S., M.O., L.S., and L.H. conducted the investigation. M.T. and R.Z performed formal analysis. U.S. performed proteomic analysis. T.Z. and N.F. developed the methodology for mass spectrometry analysis of lipids. M.T., T.Z., and N.F. performed mass spectrometry measurements and data analysis. D.Ko. performed electron microscopy. A.S. and K.G. conducted the MD simulations and computational analysis. M.T. and J.B. performed confocal microscopy and image analysis.

## Competing interests

The authors declare no competing interests.

## Acknowledgments

We thank Sarah Masser for help with the mass spec proteomics sample preparation, Gerald Rechberger for his support with the mass spectrometry investigations, Kerstin Hingerl for her support with electron microscopy experiments, and Heimo Wolinski for his help with the confocal microscopy experiments. This work was supported by the Austrian Science Fund (FWF), Field of Excellence BioHealth – University of Graz, Province of Styria, City of Graz, BioTechMed-Graz, NAWI Graz. Grant numbers: SFB Lipid Hydrolysis 10.55776/F73 (D.K., K.G., R.Z.); FWF 10.55776/P35532 (R.Z.); doc-fund “Molecular Metabolism” 10.55776/DOC50; FWF 10.55776/ESP2877024 (C.W.), FWF 10.55776/PAT3403323 (U.T.), FWF 10.55776/COE14 (U.S.). For open access purposes, the authors have applied a CC BY public copyright license to any author accepted manuscript version arising from this submission.

## Supplemental tables

**Supplementary table S1:** Lipid substrates and internal standards (IS)

| Lipid | Company | Cat# |
| --- | --- | --- |
| 18:1 PG | Avanti | 840475 |
| 18:0-18:1 PG | Avanti | 840503 |
| LPG 18:1 | Avanti | 858125 |
| Liver PI | Avanti | 840042 |
| 18:1 LPI | Avanti | 850100 |
| 18:1 PS | Avanti | 840035 |
| 18:1 LPS | Avanti | 858143 |
| 18:1 CL | Avanti | 710335 |
| 18:1 PC | Avanti | 850375 |
| 16:0-20:4 PC | Avanti | 850459 |
| 18:1 LPC | Avanti | 845875 |
| 18:1 PE | Avanti | 850725 |
| 18:1 LPE | Avanti | 846725 |
| 18:1 S,S BMP | Avanti | 857135 |
| C16-18:1 PC | Avanti | 878112 |
| C16-18:1 PE | Avanti | 878130 |
| Brain SM (Porcine) | Avanti | 860062 |
| 17:0 CE (IS) | Avanti | 700186 |
| 18:3 TAG (IS) | Larodan | 33-1830 |
| 17:0 TAG (IS) | Larodan | 33-1700 |
| 15:0 TAG (IS) | Larodan | 33-1500 |
| 17:0 DAG (IS) | Larodan | 32-1700 |
| 17:0 MAG (IS) | Larodan | 31-1700 |
| 17:0 PC (IS) | Avanti | 850360 |
| 14:0 PC (IS) | Avanti | 850345 |
| 17:0 PE (IS) | Avanti | 830756 |
| 17:0 PS (IS) | Avanti | 840028 |
| 17:1 LPC (IS) | Avanti | 855677 |
| 17:1 LPE (IS) | Avanti | 856707 |
| 17:1 LPS (IS) | Avanti | 858141 |
| 17:0 Cer (IS) | Avanti | 860517 |
| 17:0 SM (IS) | Avanti | 860585 |
| 14:0 BMP (IS) | Avanti | 857131 |
| 17:0 PG (IS) | Avanti | 830456 |
| 17:1 LPG (IS) | Avanti | 858127 |
| 14:0 Hemi BMP (IS) | Avanti | 857132 |
| Dioleoyl-1-<sup>14</sup>C PC | Hartmann Analytic | ARC0850 |
| 9,10-<sup>3</sup>H sodium oleate | Hartmann Analytic | ART0198 |

(IS): internal standard

**Supplementary table S2:**
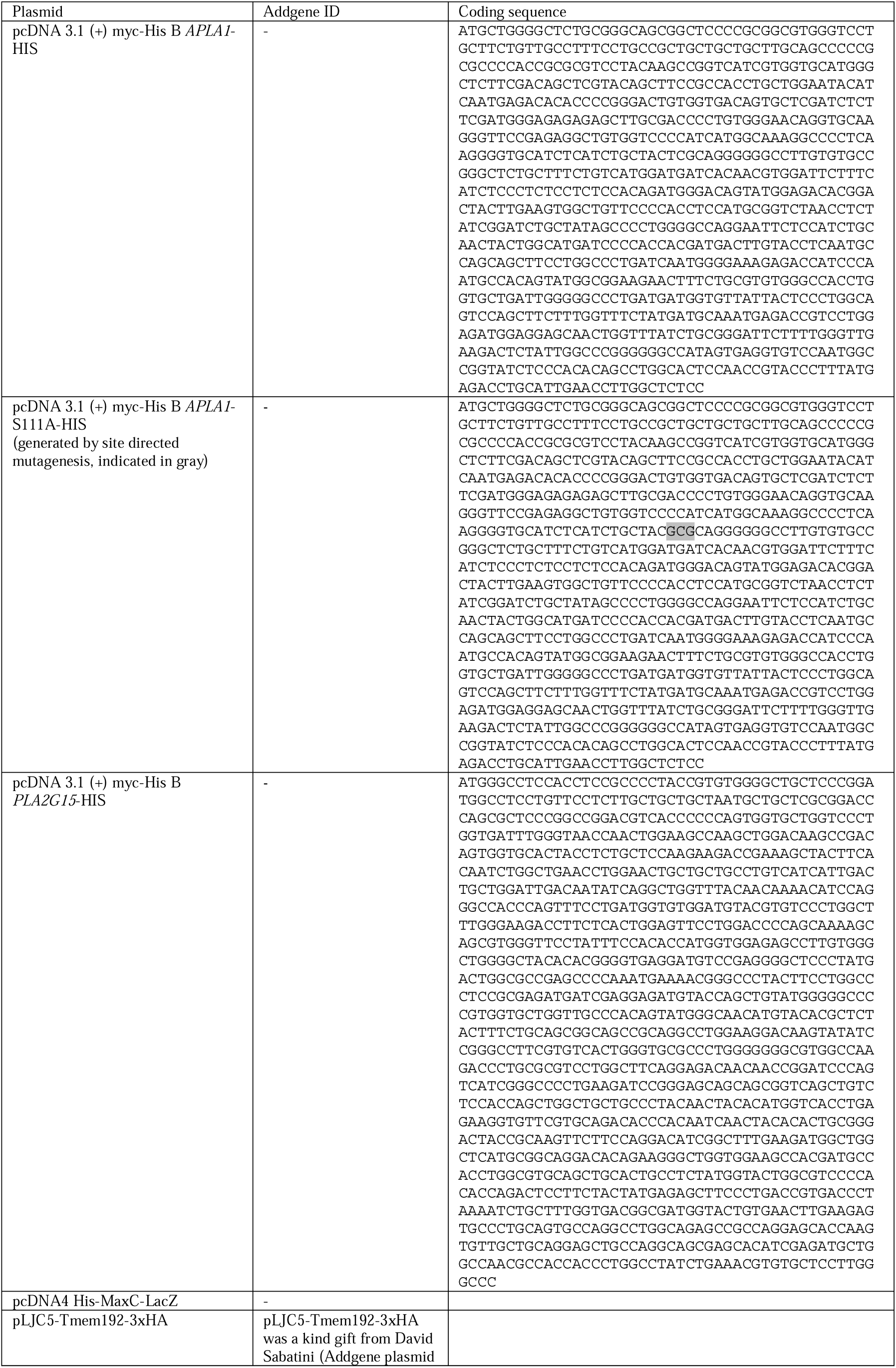
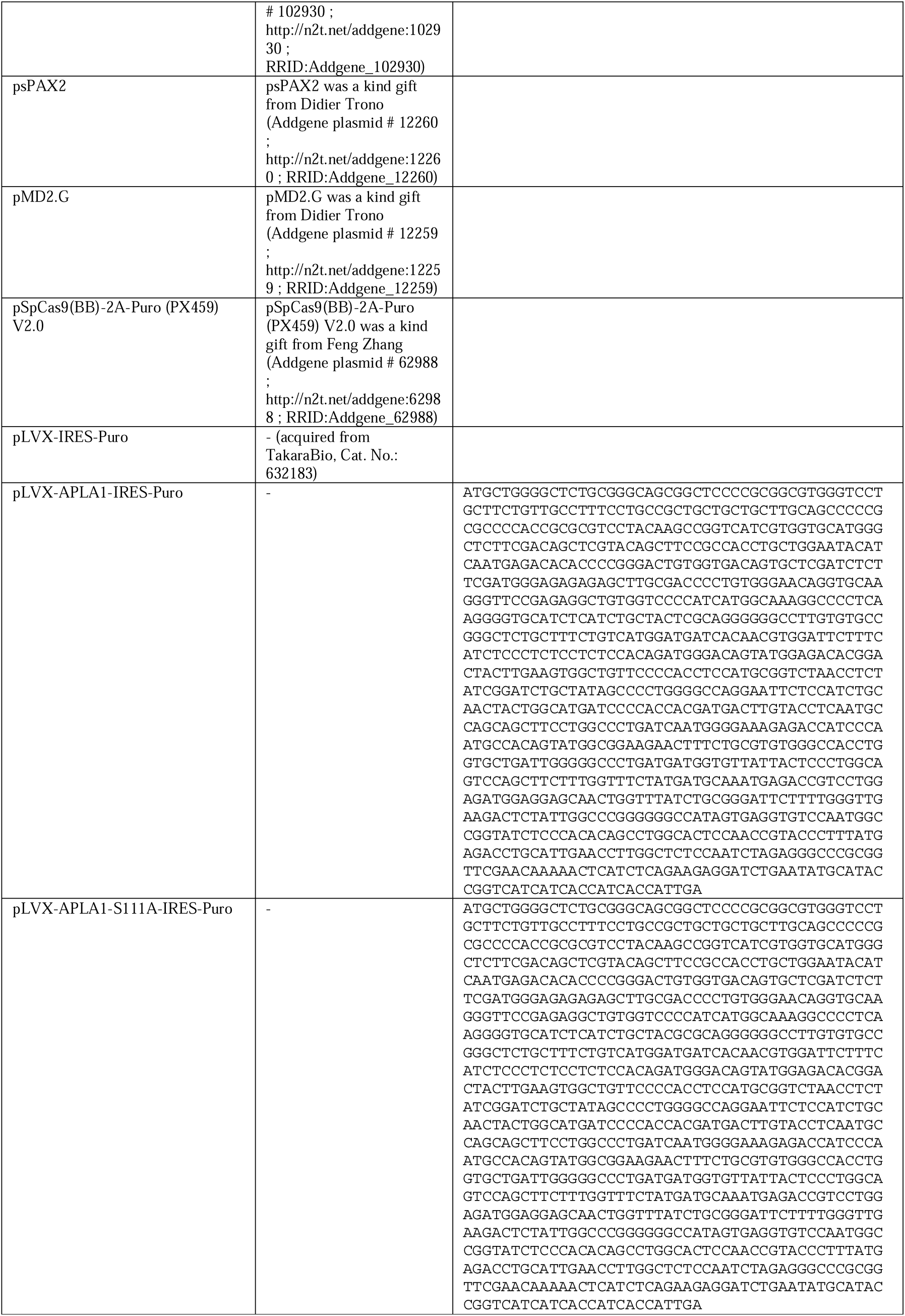
Plasmids.

**Supplementary table 3:** Antibodies.

| Antibody | Company | Cat# | Dilution* |
| --- | --- | --- | --- |
| Anti-6X HIS tag antibody | Abcam, Cambridge, UK | ab18184 | 1:3,000 |
| Rabbit Anti-GAPDH Monoclonal Antibody | Cell Signaling Technology, Danvers, USA | 2118 | 1:10,000 |
| LAMP1 (C54H11) rabbit mAb | Cell Signaling Technology, Danvers, USA | 3243 | 1:1,000 |
| Anti-LYPLA3 polyclonal Antibody (anti-PLA2G15) | Proteintech, Illinois, USA | 10863-2-AP | 1:2,000 |
| Anti-SDHA mouse mAb | Abcam, Cambridge, UK | Ab14715 | 1:1,000 |
| PDI (C81H6) Rabbit mAb | Cell Signaling Technology, Danvers, USA | 3501S | 1:1,000 |
| Goat Anti-Rabbit IgG Antibody (H+L), Peroxidase | Vector Laboratories, Newark, USA | PI-1000-1 | 1:10,000 |
| Sheep Anti-Mouse IgG ECL Antibody, HRP conjugated | Cytiva, Marlborough, USA | Na9310-1 ml | 1:10,000 |

\*all antibodies were diluted in tris-buffered saline with Tween20 supplemented with 5% blocking grade milk powder

**Supplementary table S4:** Targeted MS Transitions for BMP, PG, and LPG measurements.

| Name | Precursor Ion (m/z) | Product Ion (m/z) | Fragmentor (V) | Collision Energy (V) | Polarity |
| --- | --- | --- | --- | --- | --- |
| BMP 28:0 (14:0-14:0) (IS) | 665.4 | 227.2 | 250 | 40 | negative |
| BMP 32:0 (16:0-16:0) | 721.5 | 255.2 | 250 | 40 | negative |
| BMP 32:1 (16:0-16:1) | 719.5 | 253.2 | 250 | 40 | negative |
| BMP 34:0 (16:0-18:0) | 749.5 | 283.2 | 250 | 40 | negative |
| BMP 34:1 (16:0-18:1) | 747.5 | 281.2 | 250 | 40 | negative |
| BMP 34:2 (16:0-18:2) | 745.5 | 279.2 | 250 | 40 | negative |
| BMP 36:0 (18:0-18:0) | 777.5 | 283.2 | 250 | 40 | negative |
| BMP 36:1 (18:0-18:1) | 775.5 | 283.2 | 250 | 40 | negative |
| BMP 36:2 (18:1-18:1) | 773.5 | 281.2 | 250 | 40 | negative |
| BMP 36:3 (18:1-18:2) | 771.5 | 279.2 | 250 | 40 | negative |
| BMP 36:4 (18:2-18:2) | 769.5 | 279.2 | 250 | 40 | negative |
| BMP 36:5 (18:2-18:3) | 767.5 | 277.2 | 250 | 40 | negative |
| BMP 38:4 (18:0-20:4) | 797.5 | 303.2 | 250 | 40 | negative |
| BMP 38:5 (18:1-20:4) | 795.5 | 303.2 | 250 | 40 | negative |
| BMP 38:5 (18:2-20:3) | 795.5 | 305.2 | 250 | 40 | negative |
| BMP 38:6 (16:0-22:6) | 793.5 | 255.2 | 250 | 40 | negative |
| BMP 38:6 (18:2-20:4) | 793.5 | 303.2 | 250 | 40 | negative |
| BMP 38:7 (16:1-22:6) | 791.5 | 327.2 | 250 | 40 | negative |
| BMP 38:7 (18:3-20:4) | 791.5 | 277.2 | 250 | 40 | negative |
| BMP 40:5 (18:1-22:4) | 823.5 | 331.3 | 250 | 40 | negative |
| BMP 40:6 (18:1-22:5) | 821.5 | 329.3 | 250 | 40 | negative |
| BMP 40:6 (18:2-22:4) | 821.5 | 331.3 | 250 | 40 | negative |
| BMP 40:7 (18:1-22:6) | 819.5 | 327.2 | 250 | 40 | negative |
| BMP 40:7 (18:2-22:5) | 819.5 | 329.2 | 250 | 40 | negative |
| BMP 40:8 (18:2-22:6) | 817.5 | 303.2 | 250 | 40 | negative |
| BMP 40:8 (20:4-20:4) | 817.5 | 327.2 | 250 | 40 | negative |
| BMP 40:9 (18:3-22:6) | 815.5 | 277.3 | 250 | 40 | negative |
| BMP 42:10 (20:4-22:6) | 841.5 | 327.2 | 250 | 40 | negative |
| BMP 42:8 (20:4-22:4) | 845.5 | 303.2 | 250 | 40 | negative |
| BMP 42:9 (20:3-22:6) | 843.5 | 327.2 | 250 | 40 | negative |
| BMP 44:10 (22:4-22:6) | 869.5 | 331.2 | 250 | 40 | negative |
| BMP 44:11 (22:5-22:6) | 867.5 | 329.2 | 250 | 40 | negative |
| BMP 44:12 (22:6-22:6) | 865.5 | 327.2 | 250 | 40 | negative |
| LPG 17:1 (IS) | 495.2 | 267.2 | 180 | 28 | negative |
| LPG 14:0 | 455.2 | 227.2 | 180 | 28 | negative |
| LPG 16:0 | 483.3 | 255.2 | 180 | 28 | negative |
| LPG 16:1 | 481.3 | 253.2 | 180 | 28 | negative |
| LPG 18:0 | 511.3 | 283.2 | 180 | 28 | negative |
| LPG 18:1 | 509.3 | 281.2 | 180 | 28 | negative |
| LPG 18:2 | 507.3 | 279.2 | 180 | 28 | negative |
| LPG 20:4 | 531.3 | 303.2 | 180 | 28 | negative |
| LPG 22:6 | 555.3 | 327.2 | 180 | 28 | negative |
| PG 28:0 | 665.4 | 227.2 | 250 | 45 | negative |
| PG 32:0 (16:0-16:0) | 721.5 | 255.2 | 250 | 45 | negative |
| PG 32:1 (16:0-16:1) | 719.5 | 253.2 | 250 | 45 | negative |
| PG 34:0 (17:0-17:0) (IS) | 749.5 | 269.2 | 250 | 45 | negative |
| PG 34:1 (16:0-18:1) | 747.5 | 255.2 | 250 | 45 | negative |
| PG 34:2 (16:0-18:2) | 745.5 | 255.2 | 250 | 45 | negative |
| PG 34:2 (16:1-18:1) | 745.5 | 281.2 | 250 | 45 | negative |
| PG 34:3 (16:1-18:2) | 743.5 | 253.2 | 250 | 45 | negative |
| PG 36:0 (18:0-18:0) | 777.5 | 283.2 | 250 | 45 | negative |
| PG 36:1(18:0-18:1) | 775.5 | 281.2 | 250 | 45 | negative |
| PG 36:2 (18:1-18:1) | 773.5 | 281.3 | 250 | 45 | negative |
| PG 36:3 (18:1-18:2) | 771.5 | 281.3 | 250 | 45 | negative |
| PG 36:4 (16:0-20:4) | 769.5 | 255.2 | 250 | 45 | negative |
| PG 36:4 (18:2-18:2) | 769.5 | 279.2 | 250 | 45 | negative |
| PG 38:4 (18:0-20:4) | 797.5 | 283.2 | 250 | 45 | negative |
| PG 38:5 (18:1-20:4) | 795.5 | 281.2 | 250 | 45 | negative |
| PG 38:6 (16:0-22:6) | 793.5 | 255.2 | 250 | 45 | negative |

(IS): internal standard

**Supplementary table S5:** Targeted MS Transitions for in vitro assays.

| Name | Precursor Ion (m/z) | Product Ion (m/z) | Fragmentor (V) | Collision Energy (V) | Polarity |
| --- | --- | --- | --- | --- | --- |
| LPG 17:1 (IS) | 495.2 | 267.2 | 180 | 28 | negative |
| LPG 18:1 | 509.3 | 281.2 | 180 | 28 | negative |
| LPG 18:0 | 511.3 | 283.2 | 180 | 28 | negative |
| LPC 16:0 | 496.3 | 184.1 | 179 | 28 | positive |
| LPC 17:1 (IS) | 508.3 | 184.1 | 79 | 28 | positive |
| LPC 18:1 | 522.3 | 184.1 | 179 | 28 | positive |
| LPC 20:4 | 544.3 | 184.1 | 179 | 28 | positive |
| Cer35:1 (IS) | 552.5 | 264.2 | 106 | 28 | positive |
| Cer d18:1/16:1 | 536.5 | 264.2 | 106 | 28 | positive |
| Cer d18:1/18:0 | 566.6 | 264.2 | 106 | 28 | positive |
| Cer d18:1/22:0 | 622.6 | 264.2 | 106 | 28 | positive |
| Cer d18:1/23:0 | 636.6 | 264.2 | 106 | 28 | positive |
| Cer d18:1/24:0 | 650.6 | 264.2 | 106 | 28 | positive |
| Cer d18:1/24:1 | 648.6 | 264.2 | 106 | 28 | positive |
| DAG 34:0 (17:0-17:0) | 614.6 | 327.3 | 137 | 16 | positive |
| DAG 36:2 (18:1-18:1) | 640.6 | 339.0 | 110 | 16 | positive |

(IS): internal standard

## Notes

### Competing Interest Statement

The authors have declared no competing interest.

## References

1. Ebner, M., Fröhlich, F., and Haucke, V. (2025). Mechanisms and functions of lysosomal lipid homeostasis. Cell Chem. Biol. 32, 392–407. 10.1016/j.chembiol.2025.02.003.

2. Ludlaim, A.M., Waddington, S.N., and McKay, T.R. (2025). Unifying biology of neurodegeneration in lysosomal storage diseases. J. Inherit. Metab. Dis. 48. 10.1002/JIMD.12833.

3. Laqtom, N.N., Dong, W., Medoh, U.N., Cangelosi, A.L., Dharamdasani, V., Chan, S.H., Kunchok, T., Lewis, C.A., Heinze, I., Tang, R., et al. (2022). CLN3 is required for the clearance of glycerophosphodiesters from lysosomes. Nature 609, 1005–1011. 10.1038/S41586-022-05221-Y.

4. Shayman, J.A., and Tesmer, J.J.G. (2019). Lysosomal phospholipase A2. Biochim. Biophys. Acta Mol. Cell Biol. Lipids 1864, 932–940. 10.1016/J.BBALIP.2018.07.012.

5. Nyame, K., Hims, A., Aburous, A., Laqtom, N.N., Dong, W., Medoh, U.N., Heiby, J.C., Xiong, J., Ori, A., and Abu-Remaileh, M. (2024). Glycerophosphodiesters inhibit lysosomal phospholipid catabolism in Batten disease. Mol. Cell 84, 1354–1364.e9. 10.1016/J.MOLCEL.2024.02.006.

6. Hosios, A.M., Wilkinson, M.E., McNamara, M.C., Kalafut, K.C., Torrence, M.E., Asara, J.M., and Manning, B.D. (2022). mTORC1 regulates a lysosome-dependent adaptive shift in intracellular lipid species. Nat. Metab. 4, 1792–1811. 10.1038/s42255-022-00706-6.

7. Nyame, K., Xiong, J., Alsohybe, H.N., de Jong, A.P.H., Peña, I. V., de Miguel, R., Brummelkamp, T.R., Hartmann, G., Nijman, S.M.B., Raaben, M., et al. (2025). PLA2G15 is a BMP hydrolase and its targeting ameliorates lysosomal disease. Nature 642, 474–483. 10.1038/s41586-025-08942-y.

8. Korbelius, M., Kuentzel, K.B., Bradić, I., Vujić, N., and Kratky, D. (2023). Recent insights into lysosomal acid lipase deficiency. Trends Mol. Med. 29, 425–438. 10.1016/j.molmed.2023.03.001.

9. Breiden, B., and Sandhoff, K. (2021). Acid Sphingomyelinase, a Lysosomal and Secretory Phospholipase C, Is Key for Cellular Phospholipid Catabolism. Int. J. Mol. Sci. 22. 10.3390/IJMS22169001.

10. Kumar, M., Aguiar, M., Jessel, A., Thurberg, B.L., Underhill, L., Wong, H., George, K., Davidson, V., and Schuchman, E.H. (2024). The impact of sphingomyelin on the pathophysiology and treatment response to olipudase alfa in acid sphingomyelinase deficiency. Genetics in Medicine Open 2. 10.1016/j.gimo.2024.101888.

11. Bulfon, D., Breithofer, J., Grabner, G.F., Fawzy, N., Pirchheim, A., Wolinski, H., Kolb, D., Hartig, L., Tischitz, M., Zitta, C., et al. (2024). Functionally overlapping intra# and extralysosomal pathways promote bis(monoacylglycero)phosphate synthesis in mammalian cells. Nat. Commun. 15, 9937. 10.1038/s41467-024-54213-1.

12. Soyombo, A.A., and Hofmann, S.L. (1997). Molecular cloning and expression of palmitoyl-protein thioesterase 2 (PPT2), a homolog of lysosomal palmitoyl-protein thioesterase with a distinct substrate specificity. Journal of Biological Chemistry 272, 27456–27463. 10.1074/jbc.272.43.27456.

13. Calero, G., Gupta, P., Nonato, M.C., Tandel, S., Biehl, E.R., Hofmann, S.L., and Clardy, J. (2003). The crystal structure of palmitoyl protein thioesterase-2 (PPT2) reveals the basis for divergent substrate specificities of the two lysosomal thioesterases, PPT1 and PPT2. Journal of Biological Chemistry 278, 37957–37964. 10.1074/jbc.M301225200.

14. Gallala, H.D., and Sandhoff, K. (2011). Biological function of the cellular lipid BMP-BMP as a key activator for cholesterol sorting and membrane digestion. Neurochem Res 36, 1594–1600. 10.1007/s11064-010-0337-6.

15. Wolinski, H., and Kohlwein, S.D. (2008). Microscopic analysis of lipid droplet metabolism and dynamics in yeast. Methods Mol. Biol. 457, 151–163. 10.1007/978-1-59745-261-8_11.

16. Brown, W.J., Sullivan, T.R., and Greenspan, P. (1992). Nile red staining of lysosomal phospholipid inclusions. Histochemistry 97, 349–354. 10.1007/BF00270037.

17. Breiden, B., and Sandhoff, K. (2019). Emerging mechanisms of drug-induced phospholipidosis. Biol. Chem. 401. 10.1515/HSZ-2019-0270.

18. Laqtom, N.N., and Abu-Remaileh, M. (2026). Lysosomal Immunoprecipitation (LysoIP) from Cells and Tissues for Metabolomic and Proteomic Analyses. Methods Mol. Biol. 2976, 85–102. 10.1007/978-1-0716-4844-5_8.

19. Breiden, B., and Sandhoff, K. (2021). Acid Sphingomyelinase, a Lysosomal and Secretory Phospholipase C, Is Key for Cellular Phospholipid Catabolism. Int. J. Mol. Sci. 22. 10.3390/IJMS22169001.

20. Hamilton, J., Jones, I., Srivastava, R., and Galloway, P. (2012). A new method for the measurement of lysosomal acid lipase in dried blood spots using the inhibitor Lalistat 2. Clinica Chimica Acta 413, 1207–1210. 10.1016/j.cca.2012.03.019.

21. Chen, J., Cazenave-Gassiot, A., Xu, Y., Piroli, P., Hwang, R., DeFreitas, L., Chan, R.B., Di Paolo, G., Nandakumar, R., Wenk, M.R., et al. (2023). Lysosomal phospholipase A2 contributes to the biosynthesis of the atypical late endosome lipid bis(monoacylglycero)phosphate. Commun. Biol. 6. 10.1038/S42003-023-04573-Z.

22. Medoh, U.N., Hims, A., Chen, J.Y., Ghoochani, A., Nyame, K., Dong, W., and Abu-Remaileh, M. (2023). The Batten disease gene product CLN5 is the lysosomal bis(monoacylglycero)phosphate synthase. Science 381, 1182–1189. 10.1126/SCIENCE.ADG9288.

23. Breithofer, J., Fawzy, N., Zitta, C., Tischitz, M., Bulfon, D., Hofmann, C., Hartig, L., Wagner, C., Grabner, G.F., Pirchheim, A., et al. (2025). CLN8 enables a non-canonical phospholipid synthesis pathway. 10.64898/2025.12.18.693953.

24. Sheokand, P.K., Lacabanne, D., James, A.M., Vecchia, S. Della, Ruprecht, J.J., van der Kleij, J., Turner, K., Müller-Niva, J., Salo, M.H., Jenkins, B., et al. (2025). Stereospecific GPG acylation by CLN8 drives BMP biosynthesis and its loss leads to Batten disease. 10.64898/2025.12.18.695077.

25. Barnes, M., Raman, R., and Ekins, S. (2026). Palmitoyl-protein thioesterase-1 in health and disease. Trends Pharmacol. Sci. 47. 10.1016/j.tips.2026.01.002.

26. Gupta, P., Soyombo, A.A., Atashband, A., Wisniewski, K.E., Shelton, J.M., Richardson, J.A., Hammer, R.E., and Hofmann, S.L. (2001). Disruption of PPT1 or PPT2 causes neuronal ceroid lipofuscinosis in knockout mice. Proc. Natl. Acad. Sci. U. S. A. 98, 13566–13571. 10.1073/PNAS.251485198.

27. Gupta, P., Soyombo, A.A., Shelton, J.M., Wilkofsky, I.G., Wisniewski, K.E., Richardson, J.A., and Hofmann, S.L. (2003). Disruption of PPT2 in mice causes an unusual lysosomal storage disorder with neurovisceral features. Proc. Natl. Acad. Sci. U. S. A. 100, 12325–12330. 10.1073/PNAS.2033229100,.

28. Xu, Z., Farver, W., Kodukula, S., and Storch, J. (2008). Regulation of sterol transport between membranes and NPC2. Biochemistry 47, 11134–11143. 10.1021/bi801328u.

29. McCauliff, L.A., Langan, A., Li, R., Ilnytska, O., Bose, D., Waghalter, M., Lai, K., Kahn, P.C., and Storch, J. (2019). Intracellular cholesterol trafficking is dependent upon NPC2 interaction with lysobisphosphatidic acid. Elife 8. 10.7554/ELIFE.50832.

30. Halliwell, W.H. Cationic amphiphilic drug-induced phospholipidosis. Toxicol. Pathol. 25, 53–60. 10.1177/019262339702500111.

31. Hinkovska-Galcheva, V., Treadwell, T., Shillingford, J.M., Lee, A., Abe, A., Tesmer, J.J.G., and Shayman, J.A. (2021). Inhibition of lysosomal phospholipase A2 predicts drug-induced phospholipidosis. J. Lipid Res. 62. 10.1016/J.JLR.2021.100089.

32. Shayman, J.A., Kelly, R., Kollmeyer, J., He, Y., and Abe, A. (2011). Group XV phospholipase A2, a lysosomal phospholipase A2. Prog. Lipid Res. 50, 1–13. 10.1016/J.PLIPRES.2010.10.006.

33. Abe, A., Hinkovska-Galcheva, V., and Shayman, J.A. (2025). Assessment of bis(monoacylglycerol)phosphate isomers by thin layer chromatography using a modified solvent system. Biochem. Biophys. Rep. 44. 10.1016/j.bbrep.2025.102303.

34. Moreau, D., Vacca, F., Vossio, S., Scott, C., Colaco, A., Paz Montoya, J., Ferguson, C., Damme, M., Moniatte, M., Parton, R.G., et al. (2019). Drug-induced increase in lysobisphosphatidic acid reduces the cholesterol overload in Niemann-Pick type C cells and mice. EMBO Rep. 20. 10.15252/EMBR.201847055.

35. Tan, H.-H., Makino, A., Sudesh, K., Greimel, P., and Kobayashi, T. (2012). Spectroscopic Evidence for the Unusual Stereochemical Configuration of an Endosome-Specific Lipid. Angew. Chem. Int. Ed. 51, 533–535. 10.1002/anie.201106470.

36. Atlas Platform # AZoNano Search https://www.azonano.com/search.aspx?q=Atlas%20Platform&site=all&fsb=1.

37. Abu-Remaileh, M., Wyant, G.A., Kim, C., Laqtom, N.N., Abbasi, M., Chan, S.H., Freinkman, E., and Sabatini, D.M. (2017). Lysosomal metabolomics reveals V-ATPase and mTOR-dependent regulation of amino acid efflux from lysosomes. Science 358, 807. 10.1126/SCIENCE.AAN6298.

38. Bateman, A., Martin, M.J., Orchard, S., Magrane, M., Adesina, A., Ahmad, S., Bowler-Barnett, E.H., Bye-A-Jee, H., Carpentier, D., Denny, P., et al. (2025). UniProt: the Universal Protein Knowledgebase in 2025. Nucleic Acids Res. 53, D609–D617. 10.1093/NAR/GKAE1010.

39. Jurrus, E., Engel, D., Star, K., Monson, K., Brandi, J., Felberg, L.E., Brookes, D.H., Wilson, L., Chen, J., Liles, K., et al. (2018). Improvements to the APBS biomolecular solvation software suite. Protein Sci. 27, 112–128. 10.1002/PRO.3280.

40. Jo, S., Kim, T., Iyer, V.G., and Im, W. (2008). CHARMM-GUI: a web-based graphical user interface for CHARMM. J. Comput. Chem. 29, 1859–1865. 10.1002/JCC.20945.

41. Wu, E.L., Cheng, X., Jo, S., Rui, H., Song, K.C., Dávila-Contreras, E.M., Qi, Y., Lee, J., Monje-Galvan, V., Venable, R.M., et al. (2014). CHARMM-GUI Membrane Builder toward realistic biological membrane simulations. J. Comput. Chem. 35, 1997–2004. 10.1002/JCC.23702.

42. Huang, J., Rauscher, S., Nawrocki, G., Ran, T., Feig, M., De Groot, B.L., Grubmüller, H., and MacKerell, A.D. (2017). CHARMM36m: an improved force field for folded and intrinsically disordered proteins. Nat. Methods 14, 71–73. 10.1038/NMETH.4067.

43. Klauda, J.B., Venable, R.M., Freites, J.A., O’Connor, J.W., Tobias, D.J., Mondragon-Ramirez, C., Vorobyov, I., MacKerell, A.D., and Pastor, R.W. (2010). Update of the CHARMM all-atom additive force field for lipids: validation on six lipid types. J. Phys. Chem. B 114, 7830–7843. 10.1021/JP101759Q.

44. Jorgensen, W.L., Chandrasekhar, J., Madura, J.D., Impey, R.W., and Klein, M.L. (1983). Comparison of simple potential functions for simulating liquid water. J. Chem. Phys. 79, 926– 935. 10.1063/1.445869.

45. Lomize, M.A., Pogozheva, I.D., Joo, H., Mosberg, H.I., and Lomize, A.L. (2012). OPM database and PPM web server: resources for positioning of proteins in membranes. Nucleic Acids Res. 40. 10.1093/NAR/GKR703.

46. Essmann, U., Perera, L., Berkowitz, M.L., Darden, T., Lee, H., and Pedersen, L.G. (1995). A smooth particle mesh Ewald method. J. Chem. Phys. 103, 8577–8593. 10.1063/1.470117.

47. Hess, B., Bekker, H., Berendsen, H.J.C., and Fraaije, J.G.E.M. (1997). LINCS: A Linear Constraint Solver for Molecular Simulations. J Comput Chem 18, 14631472. 10.1002/(SICI)1096-987X(199709)18:12.

48. Bernetti, M., and Bussi, G. (2020). Pressure control using stochastic cell rescaling. J. Chem. Phys. 153. 10.1063/5.0020514.

49. Hub, J.S., De Groot, B.L., and Van Der Spoel, D. (2010). g_wham—A Free Weighted Histogram Analysis Implementation Including Robust Error and Autocorrelation Estimates. J. Chem. Theory Comput. 6, 3713–3720. 10.1021/CT100494Z.

50. Passaro, S., Corso, G., Wohlwend, J., Reveiz, M., Thaler, S., Somnath, V.R., Getz, N., Portnoi, T., Roy, J., Stark, H., et al. (2025). Boltz-2: Towards Accurate and Efficient Binding Affinity Prediction. bioRxiv. 10.1101/2025.06.14.659707.

51. Kim, A.-R., Hu, Y., Comjean, A., Rodiger, J., Mohr, S.E., and Perrimon, N. (2024). Enhanced Protein-Protein Interaction Discovery via AlphaFold-Multimer. bioRxiv. 10.1101/2024.02.19.580970.

52. Matyash, V., Liebisch, G., Kurzchalia, T. V., Shevchenko, A., and Schwudke, D. (2008). Lipid extraction by methyl-terf-butyl ether for high-throughput lipidomics. In Journal of Lipid Research (J Lipid Res), pp. 1137–1146. 10.1194/jlr.D700041-JLR200.

53. Tsugawa, H., Ikeda, K., Takahashi, M., Satoh, A., Mori, Y., Uchino, H., Okahashi, N., Yamada, Y., Tada, I., Bonini, P., et al. (2020). A lipidome atlas in MS-DIAL 4. Nat. Biotechnol. 38, 1159–1163. 10.1038/S41587-020-0531-2.

54. Kirkwood, K.I., Pratt, B.S., Shulman, N., Tamura, K., MacCoss, M.J., MacLean, B.X., and Baker, E.S. (2022). Utilizing Skyline to analyze lipidomics data containing liquid chromatography, ion mobility spectrometry and mass spectrometry dimensions. Nat. Protoc. 17, 2415–2430. 10.1038/S41596-022-00714-6.

55. White, J.B., Trim, P.J., Salagaras, T., Long, A., Psaltis, P.J., Verjans, J.W., and Snel, M.F. (2022). Equivalent Carbon Number and Interclass Retention Time Conversion Enhance Lipid Identification in Untargeted Clinical Lipidomics. Anal. Chem. 94, 3476–3484. 10.1021/ACS.ANALCHEM.1C03770.

